# Single-nucleus multi-omics identifies shared and distinct pathways in Pick’s and Alzheimer’s disease

**DOI:** 10.1101/2024.09.06.611761

**Authors:** Zechuan Shi, Sudeshna Das, Samuel Morabito, Jennifer Stocksdale, Emily Miyoshi, Shushrruth Sai Srinivasan, Nora Emerson, Arshi Shahin, Negin Rahimzadeh, Zhenkun Cao, Justine Silva, Andres A. Castaneda, Elizabeth Head, Leslie Thompson, Vivek Swarup

**Affiliations:** Department of Neurobiology and Behavior, Charlie Dunlop School of Biological Sciences, University of California, Irvine, CA 92697, USA; Institute for Memory Impairments and Neurological Disorders, University of California, Irvine, CA 92697, USA; Mathematical, Computational and Systems Biology Program, University of California, Irvine, CA 92697, USA; Department of Computer Science, University of California, Irvine, CA 92697, USA; Department of Biological Chemistry, University of California, Irvine, CA 92697, USA; Department of Pathology and Laboratory Medicine, University of California, Irvine, CA 92697, USA

## Abstract

The study of transcriptomic and epigenomic variations in neurodegenerative diseases, particularly tauopathies like Pick’s disease (PiD) and Alzheimer’s disease (AD), offers insights into their underlying regulatory mechanisms. Here, we identified critical regulatory changes driving disease progression, revealing potential therapeutic targets. Our comparative analyses uncovered disease-enriched non-coding regions and genome-wide transcription factor (TF) binding differences, linking them to target genes. Notably, we identified a distal human-gained enhancer (HGE) associated with E3 ubiquitin ligase (UBE3A), highlighting disease-specific regulatory alterations. Additionally, fine-mapping of AD risk genes uncovered loci enriched in microglial enhancers and accessible in other cell-types. Shared and distinct TF binding patterns were observed in neurons and glial cells across PiD and AD. We validated our findings using CRISPR to excise a predicted enhancer region in UBE3A and developed an interactive database, scROAD, to visualize predicted single-cell TF occupancy and regulatory networks.

**Teaser:** Comparative studies in AD and PiD reveal critical regulatory changes and identify risk gene associations for PiD.

## Introduction

Neurodegeneration is a key aspect of many neurological disorders, each with distinct molecular mechanisms and etiologies. Alzheimer’s disease (AD) is the most prevalent neurodegenerative disorder and is pathologically characterized by the progressive accumulation of amyloid-beta plaques and neurofibrillary tangles (NFTs) of tau (*1*). Conversely, Pick’s disease (PiD) is a rare behavioral variant of frontotemporal dementia (FTD) (*2, 3*), which has a prevalence of 15 to 22 per 100,000 individuals and an incidence of 2.7 to 4.1 per 100,000 individuals per year (*4*). PiD is characterized by the presence of pathological tau aggregates known as Pick bodies (*5*). Abnormal tau aggregates such as NFTs and Pick bodies alter cellular and molecular functions in the brain, but we currently do not understand the differences and similarities between these cellular changes across different tauopathies like AD and PiD (*6*).

Efforts by large-scale consortia, such as the Genetic Frontotemporal Dementia Initiative (GENFI) and ARTFL LEFFTDS Longitudinal Frontotemporal Lobar Degeneration (ALLFTD), have been instrumental in tracking and understanding brain changes before symptoms occur and in the early and moderate stages of the disease. However, PiD’s rare prevalence, combined with challenges in clinical diagnosis, has hindered comprehensive research into its pathophysiology. The scarcity of postmortem brain samples further limits our understanding of the genetic and epigenetic underpinnings of PiD. To address these challenges, comparative functional genomics analyses of different tauopathies can provide insights into shared and distinct molecular mechanisms. One recent study (*7*) has begun to explore the molecular landscapes of PiD and related tauopathies using multi-omic approaches, which have provided invaluable insights into disease-specific gene regulatory networks across brain regions like the insular cortex, motor cortex, and visual cortex. However, critical regions such as the prefrontal cortex (PFC), which is involved in higher-order cognitive functions and social behavior, remain relatively understudied.

Transcriptomic and epigenomic alterations in the PFC associated with PiD have not been extensively explored, necessitating the focus of our current study. While recent genome-wide association studies (GWAS) and fine-mapping analyses have implicated numerous genetic loci in neurodegeneration (*8–13*), much of the attention in this area is currently focused on AD over other disorders (*6*), and the functional roles of these loci are often ambiguous since they frequently reside in non-coding regions (*14–16*). The advent of single-cell epigenomics has allowed us to provide additional context for these genetic risk variants in specific cell-types (*17*), while single-cell transcriptomics has provided insights into the molecular states of NFT-bearing neurons and NFT susceptibility in AD (*18*). While these technologies have broadened our understanding of altered cellular states and gene regulatory programs in AD (*17, 19–27*), much work remains to characterize these changes in other neurodegenerative disorders and to understand their shared and unique molecular signatures.

In this study, we employed single-nucleus assay for transposase-accessible chromatin using sequencing (snATAC-seq) to characterize the open chromatin landscape and single-nucleus RNA-sequencing (snRNA-seq) to profile the gene expression of the frontal cortex in Pick’s disease donors and cognitively normal controls. We performed parallel comparative analyses of PiD datasets with our previous AD datasets to facilitate our understanding of PiD. We leveraged cell-type-specific chromatin accessibility information to model the gene-regulatory landscape of PiD and AD, identifying sets of promoter-gene links for each disease in each cell-type. We intersected these links with our internally conducted fine-mapping analyses, considering linkage disequilibrium (LD), at selected disease risk loci to nominate candidate cell-types and genes associated with non-coding risk SNPs. Further, we modeled transcription factor (TF) binding activity in each cell-type for disease and control to characterize regulatory networks and key gene-regulatory mechanisms mediated by enhancer-promoter links, allowing us to focus our attention directly on the regulators of these GWAS genes, differentially expressed genes (DEGs) and TFs. Furthermore, snRNA-seq of PiD donors corroborated some of our findings at the transcriptomic level. To validate the robustness of our insights, we highlighted a previously unknown human-gained enhancer (HGE) in excitatory neurons regulating *UBE3A*, known for its role in regulating synaptic activity, that is altered in both PiD and AD. Using CRISPR-Cas9, we excised this HGE in induced pluripotent stem cell (iPSC)-derived neurons, and we observed a subsequent downregulation of *UBE3A* using RNA-seq. Our data suggest both shared and distinct patterns of gene regulation in PiD and AD, particularly evident in the disease-enriched and specific TF activity. Furthermore, disruption in the imputed enhancer accessibility provides validation for the accurate identification of enhancer regions located more than 40kbp away from the UTR of the disease-relevant gene.

## Results

### Single-nucleus ATAC and RNA profile PiD and AD prefrontal cortex

We performed snATAC-seq on frontal cortical tissue sections of PiD and cognitively normal control cases (10x Genomics; n = 7 PiD; n = 9 control), and snRNA-seq on the same PiD and control cases (Parse Bio; n = 5 PiD; n = 3 control). Notably, our study is the first to delineate the molecular landscape within frontal cortical regions of PiD at the single-cell level. We processed our single-nucleus data separately in PiD from our previously generated snATAC-seq data of AD (10x Genomics; n = 12 late-stage AD; n = 8 control) (*17*) and snRNA-seq (10x Genomics; n = 11 late-stage AD; n = 7 control) (*17, 28*) (Fig. 1A). After quality control filtering, 83,938 snATAC-seq and 66,661 snRNA-seq profiles came from the newly generated PiD dataset (Figs. 1, B to D, S1, A and B, and Methods), and 114,784 nuclei originated from previously generated AD snATAC-seq and 57,950 nuclei were from AD snRNA-seq. In snATAC-seq, clustering analyses revealed seven major brain cell-types in this dataset - excitatory neurons (EX), inhibitory neurons (INH), astrocytes (ASC), microglia (MG), oligodendrocytes (ODC), oligodendrocyte progenitor cells (OPC), and pericytes and endothelial cells (PER-END) - annotated based on chromatin accessibility at the promoter regions of known marker genes (Fig. 1, B, D, and E). We performed label transfer using the AD dataset (*17*) as a reference and then confirmed the annotation of our excitatory and inhibitory neurons based on previously identified marker genes, namely *SYNPR* for both EX and INH neurons, *SLC17A* for EX, and *GAD2* for INH. Similarly, we annotated our glial subpopulations, including astrocyte cluster based on the *GFAP* promoter, which has been shown to increase in disease (*29*); microglia cluster based on the *CSF1R* promoter; oligodendrocyte cluster based on the *MOBP* promoter; OPC cluster based on the *PDGFRA* promoter; and PER-END cluster on the *CLDN5* promoter (Figs. 1E and S1F). Additionally, we further confirmed cell-type identities by gene activity shown in the panel of canonical cell-type marker genes (Table S1C) (*30*). In the snRNA-seq dataset, we first clustered and identified seven major brain cell-types in PiD using a panel of canonical cell-type marker genes (Figs. 1, C and D, S1, D and E, and Table S1D, Methods). These robust cell-type identifications enabled us to explore cell-type-specific alterations and molecular mechanisms underlying PiD pathogenesis with a high degree of confidence.

**Fig. 1.**
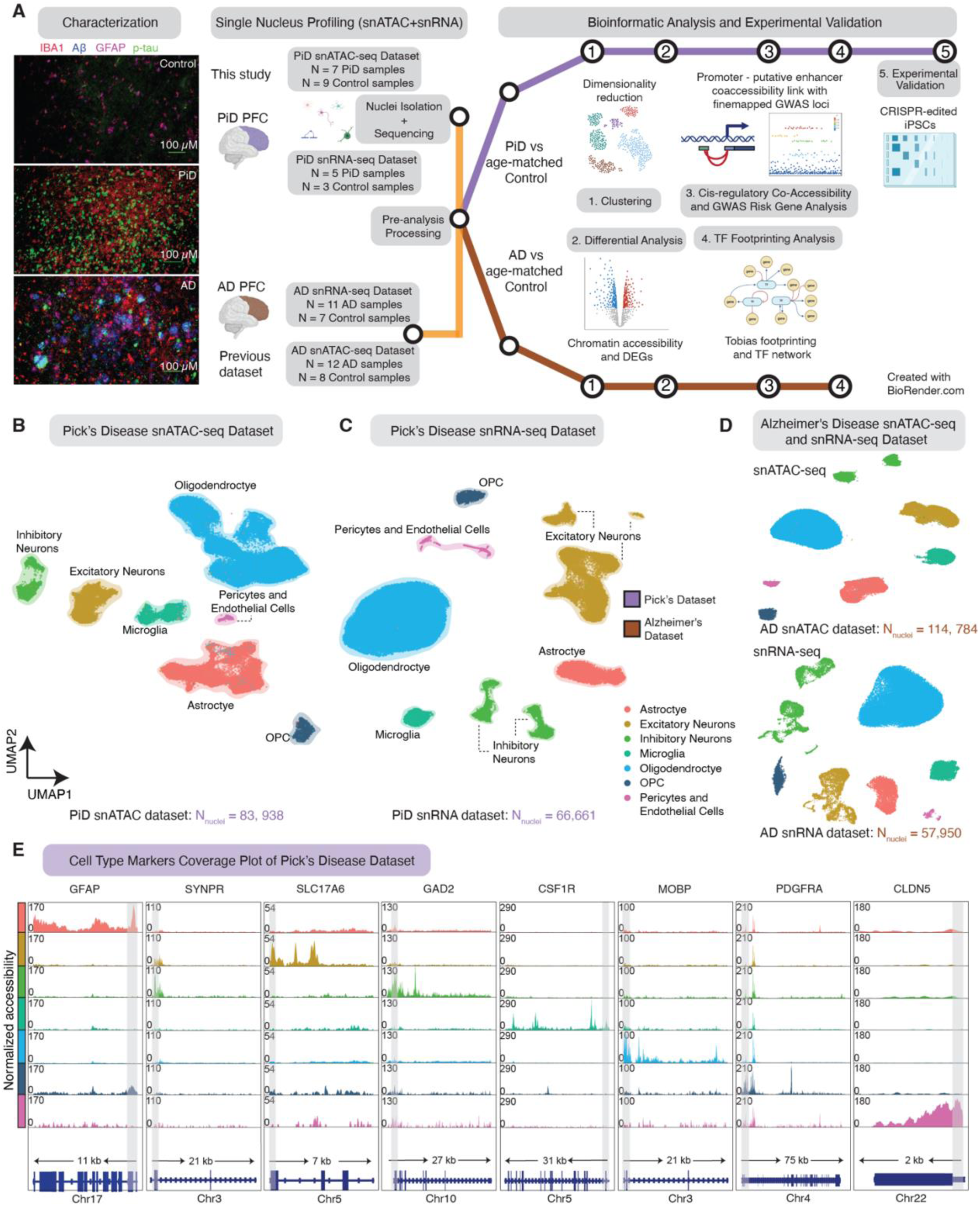
snMulti-omics for the study of cellular diversity in the PiD and AD brain. (**A**) Immunofluorescence characterization of PiD, AD and control; and schematic representation of the samples used in this study, sequencing experiments and downstream bioinformatic analyses, created with BioRender.com. Representative quadruple immunofluorescence images for IBA1 (red), GFAP (magenta), amyloid plaque (blue), and AT8/p-tau (green) from prefrontal cortex region of postmortem human brain tissues of age- and sex-matched control (n = 3), AD (n = 5) and PiD (n = 5) cases. Images were captured using Nikon ECLIPSE Ti2 inverted microscope (20X). (**B**, **C**) Uniform Manifold Approximation and Projection (UMAP) visualizations for single-nucleus ATAC-seq data (**B**) and single-nucleus RNA-seq data (**C**) from Pick’s disease and age-matched control. (**D**) UMAP visualizations for single-nucleus ATAC-seq and RNA-seq data from Alzheimer’s disease and age-matched control. (**E**) Coverage plots for canonical cell-type markers: *GFAP* (chr17:44905000-44916000) for astrocytes, *SYNPR* (chr3:63278010-63278510) for neurons, SLC17A6 (chr11:22338004-22345067) for excitatory neurons, *GAD2* (chr10:26214210-26241766) for inhibitory neurons, *CSF1R* (chr5:150056500-150087500) for microglia, *MOBP* (chr3:39467000-39488000) for oligodendrocytes, *PDGFRA* (chr4:54224871-54300000) for pericytes and endothelial cells in the PiD dataset. The gray bar within each box highlights the promoter regions.

### Promoter-enhancer linkages improve chromatin accessibility characterization

From these snATAC-seq libraries, we compiled a combined set of 609,675 reproducible peaks using ArchR (*30*) (Table S2A). To assess the robustness of our peak set, we evaluated the consistency of peak calling by comparing our data with Xiong et al.’s dataset (*27*). Overlapping peaks were defined as intersections of at least 10bp. We observed that 57% of our peaks overlapped with those reported by Xiong et al. (Fig. S2A), demonstrating a high degree of concordance. Interestingly, approximately half of these overlapping peaks intersected with more than one peak from Xiong et al.’s dataset (Fig. S2B), further highlighting the alignment of our data with previously published findings. This consistency provides confidence in the reliability of our peak set as a foundation for downstream analyses.

Building upon this robust peak set, we sought to provide functional context for non-coding distal regulatory elements with respect to cell-type and disease status. Using cis co-accessibility analyses with Cicero (*31*), we identified linkages between promoters and distal elements (Fig. 2A, Methods). Subsequently, we applied non-negative matrix factorization (NMF) to pseudobulk chromatin accessibility profiles of all distal regulatory elements linked to gene promoter regions. This analysis revealed matrix factors corresponding to epigenetic signatures of biological processes and specific cell states (Fig. 2B). We grouped CREs into discrete epigenetic modules based on the matrix factor with the highest loading for each CRE and then performed gene ontology analyses of the regulatory target genes of each module. This revealed cell-type-function-related pathways and processes regulated by these non-coding CREs, such as pathways associated with postsynaptic and synaptic activity in EX and INH, cell proliferation and migration-linked ERBB2 signaling pathway in ODC, and processes such as apoptotic cell clearance in MG. Next, using the cis co-accessibility linkages, we compared the co-accessibility strength of chromatin peak links from PiD and AD samples across the major cell lineages (Fig. 2C). These analyses revealed relatively higher correlations between PiD and AD in ODCs (Pearson R = 0.35) and ASCs (R = 0.32), with weaker correlations in other cell-types. Overall, this highlights both conserved epigenomic linkages across PiD and AD and unique regulatory landscapes specific to each condition.

**Fig. 2.**
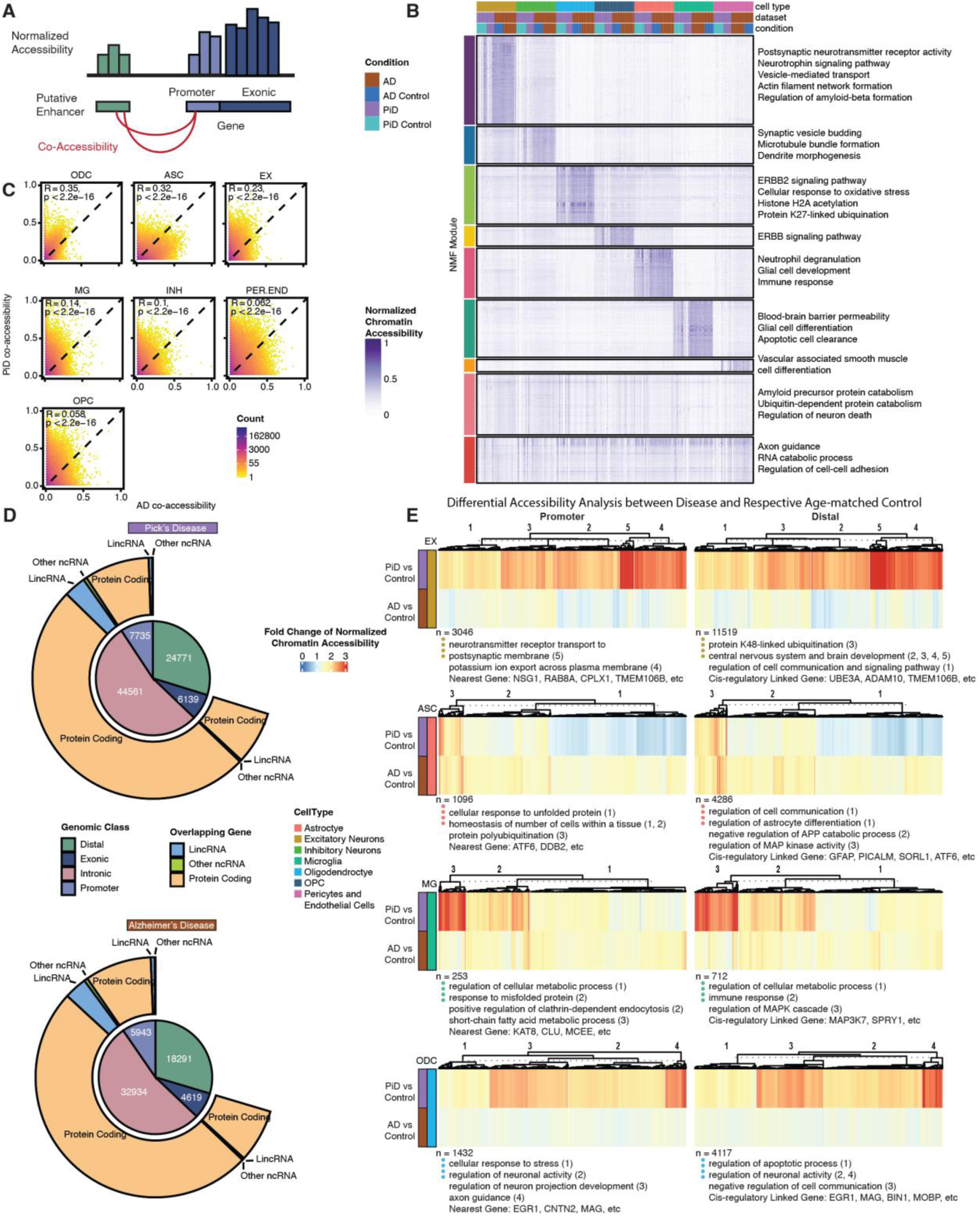
Open chromatin classification and epigenetically distinct cell-types through putative promoter-enhancer links in the human PiD and AD prefrontal cortex. (**A**) Schematics of putative promoter-enhancer linkage. (**B**) NMF heatmap of putative enhancer scaled chromatin activity in PiD, AD, and their matching controls. (**C**) Correlation heatmap of putative promoter-enhancer co-accessibility. (**D**) Peak-type and biotype classification of differentially accessible peaks (p-value < 0.05). (**E**) Heatmaps of fold changes (Disease vs. Control) on normalized chromatin accessibility of differential accessible promoters and distal in excitatory neurons, astrocytes, microglia, and oligodendrocytes (FDR-adjusted p-value < 0.05 and abs(log2FC) > 0.5), with gene ontology acquired from GREAT and examples of promoters and distal regions’ cis-regulatory linked gene as in panel (**A**).

To identify cis-regulatory elements (CREs) with altered chromatin accessibility in disease, we systematically performed differential chromatin accessibility analyses in each cell-type comparing PiD to controls and AD to controls, yielding a set of differentially accessible peaks (Table S2C). Our chromatin accessibility regions were broadly categorized by genomic features, including gene promoter, exonic, intronic, or distal regions, and we investigated these differential peaks in PiD and AD based on these categories (Fig. 2D). The majority of the differential peaks in PiD (54%) and AD (53%) were located within intronic regions. Approximately 30% of the differential peaks in both PiD and AD were distal, while 9% and 10% corresponded to promoters specifically in PiD and AD, respectively. Less than 10% of the identified differential peaks were exonic in both PiD and AD (Fig. 2D and Table S2B). These percentages were generally consistent with the peak type distribution of the entire peakset, where distal peaks comprised approximately 32%, promoter peaks made up 5%, intronic peaks constituted 55%, and exonic peaks represented 7% (Table S2A). Similar percentages were also observed in other published studies, reinforcing the robustness and consistency of our findings across different datasets (*27, 32*). Although EX was not the most sampled cell-type in PiD, our differential analyses revealed that the largest variance in the activity of CREs in the PiD dataset was observed in EX. Conversely, ASC exhibited the highest differential activity in the AD dataset (Fig. S2C).

Using this cis-regulatory linkage approach coupled with differential analyses, we used heatmaps to depict the fold changes of normalized chromatin accessibility for differentially accessible promoters, distal and intronic regions across cell-types EX, ASC, MG, and ODC (Figs. 2E, and S2E). Additionally, we incorporated gene ontology information obtained from GREAT, presenting cluster numbers alongside representative gene names. Notably, by inspecting the distal, intronic, and promoter chromatin regions and their linked regulatory target genes, we identified changes containing AD and FTD genetic risk loci, including *TMEM106B*, *ADAM10*, *SORL1*, *KAT8*, *CLU*, *BIN1* and genes involved in essential cellular activity, such as *UBE3A*. Moreover, while examining the absolute fold change of normalized chromatin accessibility, genes in EX in PiD exhibited much more robust changes than those in AD (Fig. 2E). This potentially indicates that the neuronal changes are more pronounced in PiD, likely reflecting age-associated regional differences in pathological progression, particularly in frontal cortical regions (*2, 3*). These differences align with the observation that patients with frontotemporal lobar degeneration, including PiD, often exhibit a distinct regional vulnerability (*7*) and a more rapid clinical progression compared to AD (*33*).

### Fine-mapping identifies cell-type-specific epigenomic annotations in FTD and AD

Given that the majority of variants reside in non-coding regions, around 80% of chromatin accessible peaks in distal and intronic regions (Fig. 2D), a pattern further supported by overlapped quantitative trait loci (QTLs) with chromatin accessible peaks (*34–37*) (Fig. S3D), and the limited research on disease-associated gene identification for PiD, we assert the importance of utilizing closely related FTD and AD GWAS data as reference points. Our analysis approach, an integrative method combining data from multiple modalities introduced in this study, involves overlapping snATAC-seq accessible peaks with fine-mapped GWAS SNPs, enabling us to determine whether chromosomal regions surrounding these disease-related SNPs exhibit accessibility in our dataset (Fig. S3A). However, it is important to acknowledge the inherent limitations of our study, particularly the rarity of PiD and the consequent unavailability of PiD-specific GWAS summary data with sufficient statistical power. This limitation restricts our analyses to leveraging existing knowledge and datasets to explore potential gene targets for PiD, rather than conducting direct PiD GWAS analyses. We conducted comprehensive fine-mapping, annotation, and cell-type-specific gene expression analyses, in addition to collecting publicly available predicted loss-of-function data (*38*) (gnomAD v4.0 UCSC; Methods). These efforts aimed to identify causal variants and explore the association of genetic variant-related genes with the risk of AD (*12*) and FTD (*13*) (Fig. 3). The fine-mapping analyses identified 77 lead GWAS risk SNPs with 113 credible sets, groups of genetic variants, in LD within the AD and FTD brain (posterior inclusion probability (PIP) > 0.95) overlapping with accessible peaks from the seven major cell-types. Interestingly, we found that 36 out of 113 fine-mapped causal credible sets overlapped with accessible peaks of one or two cell-types, and 16 out of 113 were present in all cell-types (Figs. 3 middle panel, and S3B, Table S3A), suggesting that some disease risk variants are relevant to a particular cell-type while others influence gene regulation across several cell-types. To expand on this, we assessed the overlap between our fine-mapped SNPs and Xiong et al.’s ROSMAP snATAC-seq cell-type-specific peaks (*27*) across seven major cell-types (Fig. 3 middle panel). In addition, we examined the uniqueness of cell-type overlaps for these fine-mapped SNPs, defined as the number of cell-types with overlapping peaks. The high proportion of overlapping peaks and the similar patterns in cell-type-specific overlaps observed in Xiong et al.’s data further support the concordance between the two datasets (Fig. S3C). To reinforce this notion, we integrated the snRNA-seq dataset from three previous studies of the AD cortex (*17, 20, 39*) and plotted the expression of genes identified from GWAS summary statistics, where each gene was associated with the lead causal SNP, from its respective control group of distinct cell-types (Fig. 3 left panel). Additionally, we corroborated the association of the lead SNP and other fine-mapped SNPs in LD with their associated genes by cross-referencing cCREs and target genes, data which can be accessed through our online interactive database, scROAD.

**Fig. 3.**
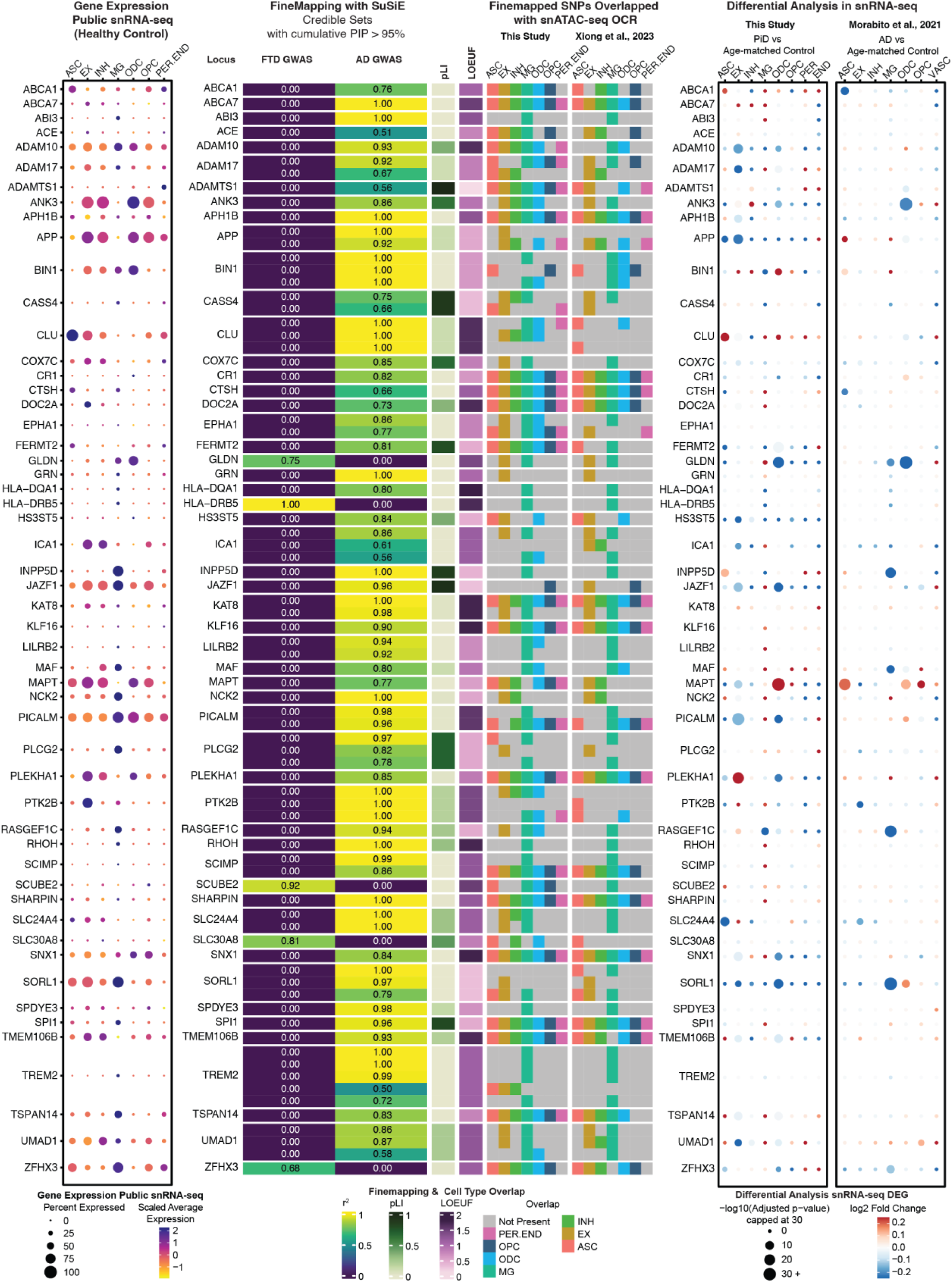
Cell-type-specific fine-mapped causal SNPs from FTD and AD GWAS risk loci. Left panel: The left dot-plot shows the gene expression in each cell-type from the control samples of three public snRNA-seq datasets (*17, 20, 39*). Middle panel: Fine-mapped SNPs from identified FTD and AD GWAS risk loci showing overlap with open chromatin regions from snATAC-seq and GWAS risk gene expression in major cell-types. The fine-mapping column using Sum of Single Effects (SuSiE) shows all of the snATAC-seq cell-type-specific open chromatin regions overlapping credible sets, defined as the groups of SNPs containing the causal variant, (PIP > 0.95). The closest gene to the credible set is indicated on the left. The r^2^ indicates the average correlation between the SNPs in the credible set. Both the probability of being loss-of-function intolerant (pLI) and loss-of-function observed/expected upper bound fraction (LOEUF) are from gnomAD (*38*) (gnomAD v4.0 UCSC). In the pLI column, a value closer to 1 indicates that the gene cannot tolerate protein-truncating variation. In the LOEUF column, a value closer to 0 indicates that the GWAS risk gene is constrained or mutation intolerant. The overlapped snATAC-seq OCR columns, including SNPs overlapped with peaks in this study and in Xiong et al. (*27*), reflect the cell-types of those causal SNPs from a credible set that are present or absent. Right panel: The two dot-plots on the right show the snRNA-seq differentially expressed GWAS genes in each cell-type between PiD and age-matched control samples, and between AD and age-matched control samples (17). A complete set of the fine-mapped SNPs and credible sets with a PIP > 0.95 shown for FTD and AD is available in Table S3. Data on fine-mapped SNPs with cCREs and their associated target genes can be accessed through our online interactive database, scROAD.

For AD, our analyses revealed that more than half of the 113 fine-mapped signals overlapped with accessible peaks found in microglia, a cell-type of particular interest in AD research. Notably, these peaks encompassed several known AD GWAS genes that have been extensively studied in microglia, including *ABCA1*, *ADAM10*, *ADAM17*, *BIN1*, *INPP5D*, *NCK2*, *PICALM*, and *TREM2* (Fig. 3 middle panel). The enrichment of GWAS risk signals within microglia was consistent with the established pathophysiological role of these cells, particularly their involvement in inflammation in AD (*40*). AD risk variants at the *INPP5D* locus were found in accessible chromatin regions exclusively in microglia, and the *INPP5D* gene was expressed almost specifically in microglia as well (Fig. 3 middle panel).

While previous studies have demonstrated the enrichment of AD genetic risk SNPs specifically in microglia (*17*), we note that these risk genes are expressed in several cell-types. For example, the risk variants of *ADAM10* overlapped with accessible peaks from EX, INH, MG and ODC and its gene expression was detected across all cell-types. As the major constituent of α-secretase, *ADAM10* cleaves APP towards a non-amyloidogenic pathway, thereby preventing Aβ generation (*41*). Furthermore, fine-mapping analyses revealed that *BIN1* risk variants, a major risk factor for AD known to induce tau- and isoform-dependent neurotoxicity (*42, 43*), predominantly localize to accessible peaks associated with ASC, MG, ODC, and OPC. These findings give credence to previously reported disparate findings on the effects of *BIN1* SNPs in microglia (*44*) and oligodendrocytes (*17*). Considering that a given gene can often be expressed in multiple cell-types, it is crucial to exercise caution when analyzing the effects of variants, as these effects may vary greatly among different cell-types. Similarly, GWAS variants in the *TREM2* gene were identified within accessible peaks primarily associated with MG and EX. *TREM2* plays a crucial role in various cellular processes, including cell proliferation, survival, phagocytosis, and regulation of inflammation (*45*). Notably, its defensive response against AD pathology, coupled with its upregulation in reactive microglia surrounding amyloid plaques, has been consistently observed across multiple studies, both in mouse models and human samples (*40, 46, 47*).

Complementing our analyses of AD risk loci, we also performed fine-mapping analyses on GWAS risk loci for FTD, aiming to propose possible risk genes for FTD subtype PiD (Fig. 3, and Table S3A and B), with five of them intersecting with accessible chromatin regions in our snATAC-seq dataset. For example, one of the fine-mapped FTD risk loci, *SLC30A8*, encodes a zinc transporter and is a susceptible GWAS locus for type 2 diabetes (*48*). Strikingly, there is a notable increase in the prevalence of both type 2 diabetes and dementia in older adults (*49*). We speculate that *SLC30A8* could be an indirectly related risk locus for FTD. Moreover, among the identified FTD risk loci, *GLDN* stands out as another intriguing candidate. *GLDN* encodes gliomedin, a crucial protein involved in the formation of the nodes of Ranvier (*50*). These nodes are critical structures along the neural axons where action potentials are regenerated. Disruption of the nodes of Ranvier can result in the failure of the electrically resistive seal between the myelin and the axon, ultimately contributing to various neurological diseases (*51*). Given the fundamental role of gliomedin in maintaining axonal integrity, investigating *GLDN* variants within specific cell-types may provide valuable insights into their potential involvement in FTD pathogenesis. In particular, our snRNA-seq differential analyses between PiD and age-matched controls revealed that *GLDN* was statistically significantly downregulated (Fig. 3 right panel). Besides *GLDN*, in our snRNA-seq analyses, some of the AD GWAS genes, such as *ADAM10*, *ADAM17*, *BIN1*, *APP*, *CLU*, *JAZF1*, *MAPT*, *PICALM*, *PLEKHA1*, *SLC24A4*, *SORL1*, and *UMAD1*, were also differentially expressed in PiD (Table S4 A and B). While risk loci have been identified in our GWAS studies and cis-regulatory-linked risk genes, several open chromatin regions that overlap with AD/FTD GWAS SNPs (Fig. 3) are also differentially accessible regions (DARs) in the PiD or AD vs. their age-matched control comparison. There are 41 EX DARs and 12 ODC DARs in the PiD dataset and 21 DARs in the AD dataset, of which some overlap with SNPs identified in the AD/FTD GWAS fine mapping (Fig. S3E). These findings underscore the shared genetic mechanisms across tauopathies while also reflecting cell-type-specific chromatin accessibility differences. Furthermore, AD GWAS genes show a strong overlap with differentially expressed genes in PiD (Fisher’s Exact Test: p-value < 2.2×10^−16^, Table S3C), suggesting that these associations are not random. However, it remains crucial to determine how fine-mapped signals specifically relate to PiD. By integrating these genetic findings with our multi-omics data, we can gain deeper insights into the complex interplay between genetic risk factors and cellular processes contributing to PiD and AD pathology, particularly with regard to regulatory non-coding regions and gene expression in the corresponding cell-types.

### Neuron TF binding occupancy reveals dysregulation in PiD and AD

To uncover gene regulatory mechanisms impacting neurons and glial cells in PiD and AD, we investigated co-accessible enhancer-promoter regions, focusing on genome-wide and gene-specific TF differential binding activities. Various gene regulatory network (GRN) approaches (*52, 53*) based on motif enrichment analyses often infer TF activity from overrepresented motifs without distinguishing functional binding from chromatin relaxation (*24, 54*), resulting in false positives (*55*), where “relaxed” open chromatin regions may not always indicate meaningful regulatory activity (Fig. S4 A and B). To address this limitation, our approach incorporates TF footprinting using the TOBIAS tool (*56*) to directly measure TF occupancy. This strategy resolves functional from non-functional motif occurrences by identifying functional enhancer-TF interactions and advances our understanding of FTD and AD genetic risk signals, including the role of fine-mapped SNPs in putative regulatory functions. We performed chromatin cis co-accessibility and TF occupancy prediction analyses on 609,675 cCREs (Table S2A) to examine disease-enriched signals in both PiD and AD. For each predominant cell-type, we implemented cis-regulatory co-accessibility (*31*) and trans-regulatory occupancy prediction (*56*), dividing the cells into PiD, AD, and their corresponding controls for detailed examination (Fig. 4A).

**Fig. 4.**
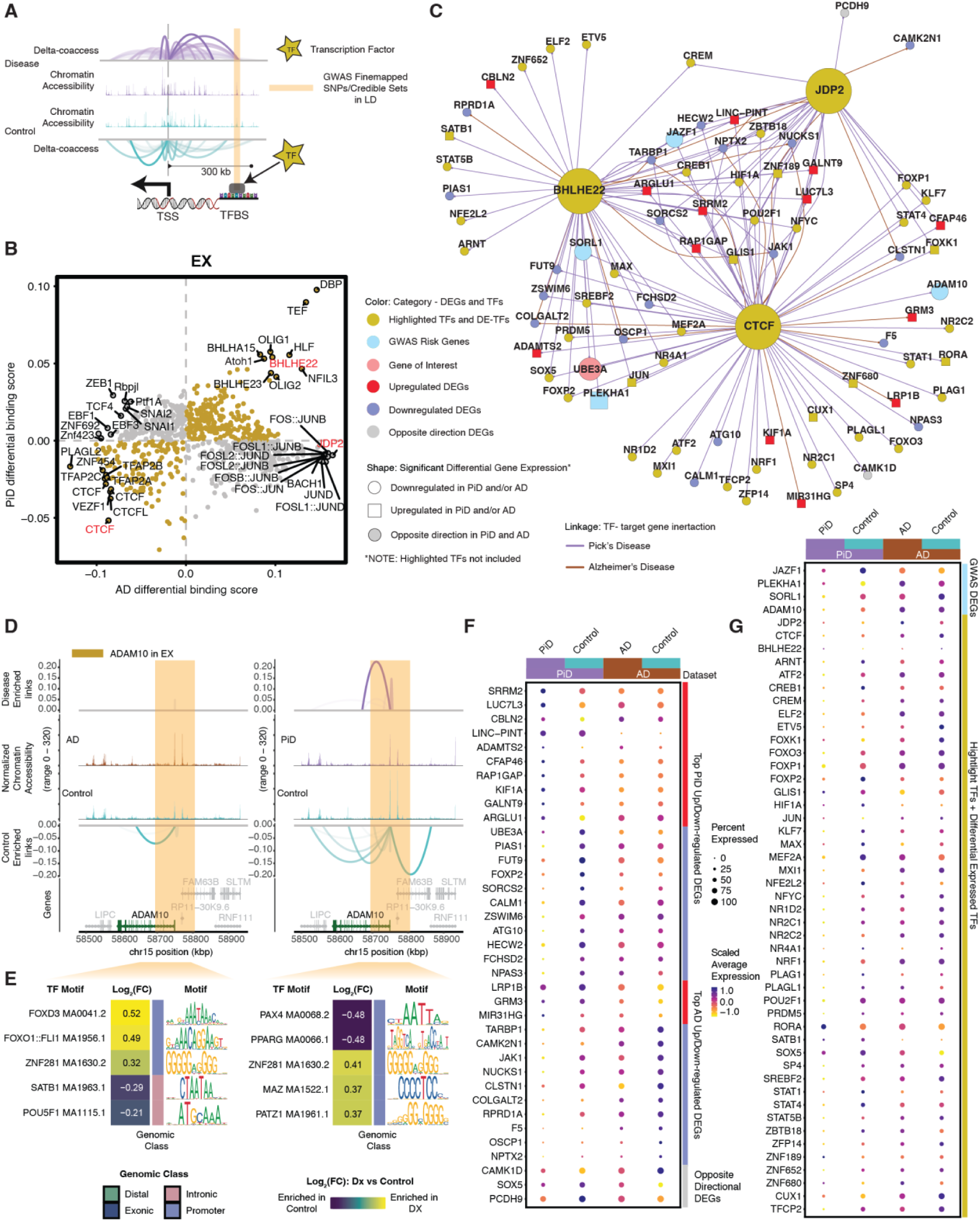
Excitatory neuronal related transcription factor dysregulation and gene expression changes associated with PiD and AD pathology. (**A**) Schematic of co-accessible mapping between putative enhancer and promoter for the target gene as well as the TF binding activity at its local regions. (**B**) Genome-wide Tn5 bias-subtracted TF differential footprinting binding scores of PiD and AD in excitatory neurons (EX) compared to the corresponding controls. (**C**) Transcription factor (TF) regulatory networks showing the predicted candidate target genes for the following TFs: *CTCF*, *BHLHE22*, and *JDP2* in EX. Highlighted transcription factors and other differentially expressed TFs are shown in yellow. Upregulated differentially expressed genes are shown in red and square. Downregulated differentially expressed genes are shown in blue and in a circle. The gene of interest, *UBE3A*, is downregulated, shown in pink and in a circle. Differentially expressed GWAS risk genes are displayed in bright blue. Edges representing the linkage of TF-target gene regulation are shown in purple for PiD and sienna for AD. (**D**) Delta co-accessibility of *ADAM10* and its open chromatin regions in EX for both AD and PiD with their corresponding controls. Highlighted regions in dark yellow represent all SuSiE fine-mapped SNPs (Fig. 3) close to the target gene. (**E**) Fold changes of TFs binding in the SuSiE fine-mapped regions for both AD and PiD. (**F**) Dot-plot of differentially expressed genes in PiD and AD versus their respective controls. (**G**) Dot-plot of differentially expressed GWAS risk genes and TFs in PiD and AD versus their respective controls.

Our integrated cis- and trans-regulatory analyses approach allows us to explore disease-enriched enhancer-promoter links, TF differential binding activity, and motif binding site disruption in neurons. Genome-wide TF differential binding scores were calculated in PiD and AD with their matching controls in neurons (Fig. 4B). We identified transcription factors *BHLHE22*, a TF previously indicated to play a key role in neural cell fate (*57*), along with other TFs (p-value < 0.05), which exhibit shared and enhanced binding activity in both PiD and AD compared with their respective controls. *JDP2*, a TF involved in apoptosis (*58*), along with other TFs, demonstrates increased binding activity only in AD. *CTCF*, a transcriptional regulator that acts on enhancers, promoters, and gene bodies (*59*), together with other TFs in the lower left quadrant, displays decreased binding activity in both PiD and AD compared to their controls. To ensure that the observed changes were not biased toward surviving neurons or influenced by sampling quality control, we calculated a neuronal vulnerability module score based on a list of vulnerable genes associated with reduced expression in disease conditions collected from Mathys et al. (*60*). This analysis demonstrated that our PiD snRNA-seq data includes both surviving and vulnerable neurons, with vulnerability module scores higher in controls compared to PiD (Fig. S4C).

We investigated the binding of select transcription factors to the enhancer regions of their target genes in neurons to contextualize their variable binding activity. (Fig. 4 C to E). To accomplish this, a gene regulatory network for the transcription factors *BHLHE22*, *CTCF*, and *JDP2* was established in both PiD and AD datasets for excitatory neurons (Figs. 4 B and C, and S4D). Several genes implicated in AD GWAS, including *JAZF1*, *SORL1*, *PLEKHA1*, and *ADAM10*, exhibited differential expression in EX in individuals with PiD, providing possible insights into shared molecular mechanisms between PiD and AD, suggesting potential convergent pathways underlying neurodegeneration in these conditions (Fig. 4C). Importantly, some of these fine-mapped GWAS genes within the TF network were further supported by significant eQTLs (*36*) (p-value < 1×10^−5^) observed in EX, reinforcing their importance in this cell-type (Fig. S4E). The differentially expressed TFs and genes we identified, positioned in the center of the network, are under the regulation of all three highlighted factors: *CTCF*, *JDP2*, and *BHLHE22*. Those regulated by two or a single TF are depicted on the outer ring of the network. We stress that these findings merely represent a simplified depiction of a highly complex regulatory network. Gene targets within this network are acknowledged to be subject to regulation, but it is important to note that the highlighted transcription factors do not solely govern their regulation.

To complement our analyses of TF trans-regulatory network in neurons, we aimed to discern cis-regulatory elements and DNA-binding motifs that are enriched in either disease or control conditions, particularly within regions containing fine-mapped SNPs. Through the integration of the co-accessibility map with chromatin accessibility signals and GWAS statistics across the genomic axis, we elucidated potential disruptions in cis-regulatory relationships caused by causal disease variants in a GWAS gene, *ADAM10*, which is also differentially expressed (Fig. 4 D and E). Additionally, we conducted sequence analyses to identify motifs that are disrupted in comparison to control conditions. This procedure was executed with the aim of assessing disease or control gene local enhancer accessibility and predicting potential disruptions in TF binding.

We found alterations in the cis-regulatory mechanisms of *ADAM10* in AD, a prominent anti-amyloidogenic candidate gene in AD pathology (*41*) (Fig. 4D). These changes were identified in proximity to the fine-mapped lead *ADAM10* SNP, rs442495, and in its strong LD block, potentially disrupting the DNA-binding motif. Consequently, these disruptions may result in diminished transcription factor (TF) binding activity in disease compared to their corresponding control group. We further investigated the gene locus TF binding activity in those highlighted fine-mapped accessible regions. We selected five TFs from the top-ranked TFs based on the average log2(fold change) of the TF binding score. For example, we found forkhead box O1 (*FOXO1*), SATB homeobox 1 (*SATB1*), *POU* class 5 homeobox 1 (*POU5F1*), Paired box 4 (*PAX4*), and peroxisome proliferator-activated receptor (*PPAR*) transcription factors enriched in highlighted regions identified for *ADAM10* in EX (Fig. 4e). Previous studies have investigated the potential roles of *FOXO1*, *SATB1*, and *POU5F1* in the development of AD (*61–63*). Notably, *FOXO* TF families were indicated as mediators of stress adaptation, which promotes the resilience of cells as a key regulator in other pathways, such as metabolism, cell cycle, and redox regulation (*64*). The transcription factor *PAX4* has been investigated in the contexts of both AD and Type 2 Diabetes (*T2D*), and is known to function as a key link in the common pathways of both diseases (*65*).

To thoroughly examine the differences in gene expression in EX between disease and control groups, we arranged and compared all selected differentially expressed genes (DEGs) and transcription factors (TFs) side by side for PiD and AD (Fig. 4 F and G). The fold change in gene expression indicates the robustness of biological changes between diseases and highlights the role of certain genes and TFs in disease development. Among those top-selected genes, identified based on its absolute fold change and cis-regulatory co-accessibility score, *CALM1* has been linked to the progression from mild cognitive impairment (MCI) to AD through involvement in the neurotrophin signaling pathway, which contributes to neuronal development, survival, and plasticity (*66*). Additionally, *CALM1* participates in dysregulated ligand-receptor (LR) interactions (*67*). Its downregulation in both PiD (FDR-adjusted p-value = 5.57×10^−6^, Table S4A) and AD (FDR-adjusted p-value = 0.011, Table S4B) samples suggests a common role of *CALM1* in the pathogenesis of both diseases. Similarly, *TARBP1* showed a notable decrease in both PiD and AD (Fig. 4 C and F). *TARBP1* encodes the TAR RNA binding protein 1 (TRBP), which participates as a methyltransferase enzyme in post-transcriptional gene regulation through its involvement in RNA processing pathways and is associated with inattention symptoms (*68*). Whereas we had previously identified the differential regulation of the distal enhancer of the *UBE3A* gene (Fig. 2E), we further found that *UBE3A* expression was statistically significant decreased in EX in PiD (FDR-adjusted p-value = 1.04 ×10^−14^, Table S4A), regulated by *CTCF* and *BHLHE22* (Fig. 4 C and F).

We conducted a detailed examination of the alterations in TFs’ expression levels between diseased and control states. Our analyses revealed a general trend of pervasive downregulation of TF expression across PiD samples, when compared to the changes observed between AD and its respective control group, despite a few TFs showing upregulation (Figs. 4G, and S4 F and G). A similar trend was observed in Rexach et al.’s behavioral variant FTD dataset (*7*), where reduced TF expression was consistent across disease samples compared to controls (Fig. S4H). This trend highlights a broader downregulation of TFs in PiD but not in AD (Fig. S4G). These unique regulatory patterns displayed in PiD emphasize the complexity of these mechanisms. Among the differentially expressed TFs, we observed *RORA*, which plays an essential role in energy and lipid metabolism (*69*), is statistically significantly upregulated in both PiD (FDR-adjusted p-value = 1.14 ×10^−20^, Table S4A) and AD (FDR-adjusted p-value = 0.002, Table S4B). Aberrant energy metabolism is the critical factor for cell integrity maintenance and neurodegeneration. Another notable differentially expressed TF, *STAT1*, demonstrated a different expression pattern across PiD (FDR-adjusted p-value = 4.61 ×10^−14^, Table S4A) and AD (not statistically significant), implying its distinct involvement in the regulatory mechanisms underlying different neurodegenerative disorders or different stages of disorders. Prior research has indicated that decreased *STAT1* expression correlates with a higher risk of conversion to MCI and can be considered a preclinical indication of AD development (*70*). The preceding analyses and these data provide a likely genetic mechanism for two distinct dementias, based on differential TF binding activity on the enhancer or promoter regions of its target gene, coupled with analyses shown on gene expression.

In inhibitory neurons (INH), we highlighted two TFs, *JDP2* and *NRF1*. *JDP2*, a transcription factor linked to apoptosis (*58*), emerged as one of the top TFs based on Tn5 bias-subtracted TF differential footprinting binding scores in AD. *NRF1*, a master regulator of proteasome genes, plays a critical role in proteasome-mediated protein degradation, a process whose dysregulation has been implicated in neurodegenerative diseases (*71*). In contrast to *JDP2*, although *NRF1* was not among the top TFs with differential footprinting binding scores in excitatory neurons (EX), it was identified as one of the top factors within the INH TF network (Figs. S5 A to C). Notably, both *JDP2* and *NRF1* are also expressed in EX, suggesting shared regulatory mechanisms between these neuronal subtypes (Figs. 4 B and G, and S4 F and G). These findings complement our results in EX, highlighting both cell-type-specific and shared transcriptional regulatory mechanisms in neurons, which may have important implications for understanding their roles in neurodegeneration.

### TF binding occupancy reveals glial responses in PiD and AD

We investigated the regulatory role of several TFs in glial cells in PiD and AD. Given the importance of TFs in modulating gene expression, we focused on identifying the top differential binding TFs, distinguishing those specific to PiD and those shared with AD. Among the selected TFs, we explored the regulatory effects of microglial TF *SPI1*, a well-known AD GWAS risk gene (*12*), Friend leukemia integration 1 (*FLI1*), and Transcription Factor Dp-1 (*TFDP1*) (Figs. 5A, and S6D), to shed light on the potential roles of these TFs in the pathogenesis of PiD and AD. In our snATAC-seq analyses of microglial cells, we observed increased differential binding activities of *FLI1* and *SPI1* in both PiD and AD. *SPI1* is known to be associated with the normal development of microglial cells in the brain (*72*), and Ets-related transcription factor *FLI1* has been established as a regulator of gene activity during cellular differentiation (*73*) (Fig. 5A). However, *TFDP1*, a potential global modulator of chromatin accessibility by controlling histone transcription (*74*), shows contrasting differential binding activities when comparing PiD with AD (Fig. 5A), suggesting potential discrepancy in genome-wide *TFDP1* TF binding activity between diseases. Among the top-selected targets, we observed a statistically significant downregulation of *MAF* in AD (FDR-adjusted p-value = 2.28×10^−8^, Table S4B), a gene identified as an AD GWAS risk gene and a differentially expressed TF (12), regulated by *SPI1*. Interestingly, we did not observe any notable difference in *MAF* expression in PiD (Figs. 5B, and S6A). Additionally, another AD DEG, *CX3CR1* (FDR-adjusted p-value = 1.53×10^−25^, Table S4B), was also regulated by SPI1 but not markedly dysregulated in PiD. *CX3CR1* has been implicated in both neuroprotective and detrimental effects by regulating inflammation in neurological disorders (*75*). Furthermore, our analyses revealed the differential expression of several other GWAS risk genes regulated by *TFDP1*, *FLI1*, and *SPI1*, including *GLDN*, *ZFHX3*, *USP6NL*, *SORL1*, *MS4A4A*, *INPP5D*, and *RASGEF1C* (*12*).

**Fig. 5.**
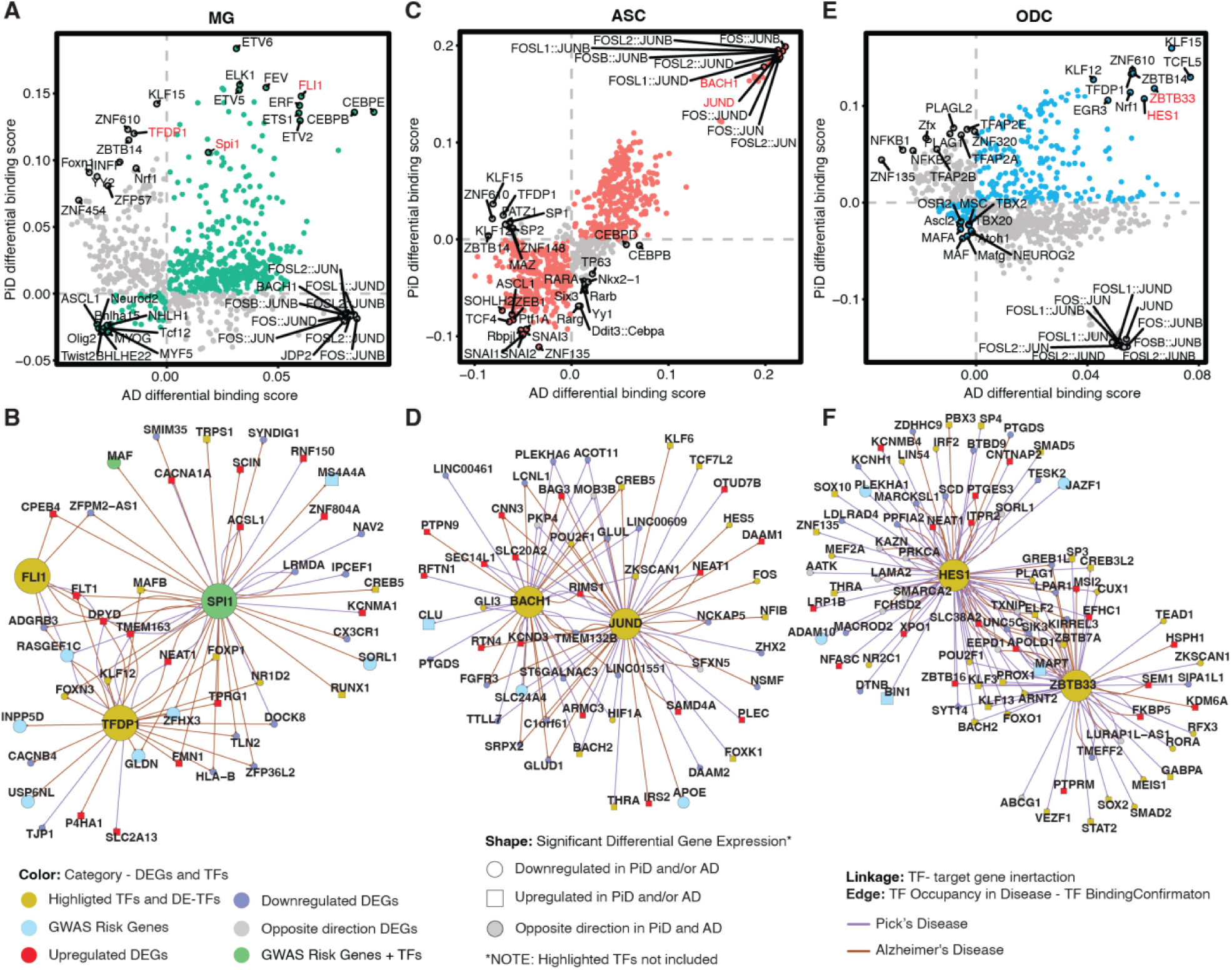
Glial changes in transcription factor dysregulation and gene expression in PiD and AD progression. (**A**, **C**, **E**) Genome-wide Tn5 bias-subtracted TF differential footprinting binding score of PiD and AD in microglia (MG) (**A**), astrocytes (ASC) (**C**), and oligodendrocytes (ODC) (**E**) compared to their corresponding controls. (**B**, **D**, **F**) TF regulatory networks showing the predicted candidate target genes for MG (**B**), ASC (**D**), and ODC (**F**). Highlighted transcription factors and differentially expressed TFs are shown in yellow. Upregulated differentially expressed genes are shown in red and square. Downregulated differentially expressed genes are shown in blue and in a circle. The differentially expressed GWAS risk genes are displayed in bright blue. Edges representing the linkage of TF-target gene regulation are shown in purple for PiD and sienna for AD.

In astrocytes, we observed a consistent trend among most TFs, where the majority displayed either increased or decreased binding scores in both PiD and AD. Notably, a subgroup of TFs from the activating protein-1 (AP-1) family, namely *JUND*, *JUNB*, and *FOS*, exhibited pronounced enrichment in both PiD and AD (Figs. 5C, and S6E). For instance, *JUND* from the AP-1 TF family, known for its strong correlations with pTau and amyloid-beta (*76*), demonstrated similar patterns. Additionally, *BACH1*, primarily recognized as a transcriptional suppressor (*77*), showed a positive correlation with both PiD and AD. These findings suggest some potential convergence of top-selected TFs’ activity in astrocytes across PiD and AD. Specifically, *JUND*’s inferred role in astrocyte *APOE* expression, which is shown to be downregulated in AD (FDR-adjusted p-value = 3.14×10^−4^, Table S4B) but not statistically significant in PiD (Figs. 5D, and S6B), underscores its involvement in AD-related processes. At the same time, we identified hypoxia-inducible factor-1 alpha (*HIF1A*), regulated by both *JUND* and *BACH1*, as downregulated in PiD (FDR-adjusted p-value = 4.35×10^−6^, Table S4A) but not statistically significant in AD, which may align with previous reports suggesting that the loss of *HIF1A* within astrocytes protects neurons from cell death (*78*). Our observations underscore potential regulatory changes in astrocytes, characterized by the regulatory activation mediated by AP-1 family TFs and the transcriptional suppression facilitated by *BACH1*. Furthermore, the dysregulation of *APOE* expression and *HIF1A* levels in astrocytes highlights the complex regulatory networks that influence astrocyte function and contribute to disease progression in AD and PiD.

In oligodendrocytes, we observed a predominant trend where the majority of TFs exhibited either increased binding activity in both PiD and AD or unique patterns specific to each disease state (Figs. 5E, and S6F). Noteworthy among these are the transcriptional suppressors *HES1* and *ZBTB33* (*79, 80*), which displayed enriched differential binding scores in both PiD and AD. Moreover, our analyses revealed that these two transcriptional repressors were associated with the downregulation of *ADAM10*, *PLEKHA1*, and *JAZF1*, and the upregulation of *BIN1* and *MAPT*, consistent with broader transcriptional changes across multiple DEGs (Figs. 5E, and S6C). This suggests the intricate and multifaceted nature of the transcriptional processes, which may be relevant to both PiD and AD, or specific to one of these conditions, indicating shared or condition-specific regulatory mechanisms. Furthermore, *MAPT*, a gene encoding tau protein to keep the function of microtubules and axonal transport, which *ZBTB33* also regulates, is differentially expressed in both PiD and AD. Additionally, the downregulation of *FOXO1*, known to protect against age-progressive axonal degeneration (*81*), further underscores the intricate interplay between transcriptional regulation and neurodegenerative processes in oligodendrocytes.

To further evaluate the reliability of the observed transcriptional changes, we analyzed the percentage of cells expressing selected genes, grouped by samples and color-coded by diagnosis (Fig. S7 A to C). This analysis encompassed both downregulated and upregulated DEGs in PiD, as well as GWAS risk genes expressed in the selected cell-type, as highlighted in TF regulatory network (Figs. 5 B, D, and F). Despite the inherent sparsity of snRNA-seq data, the percentage expression of genes exhibited consistent patterns across individuals, with no single sample disproportionately influencing the results. These stable trends across libraries and individuals affirm the robustness of the observed differences in gene expression and support the conclusions drawn from these analyses.

### Cis-regulatory linked HGE impacts gene expression in disease synaptic pathology

We have elucidated the shared and distinct changes in the pathways between these two frontal cortical degenerative diseases related to the prominent features, glial activation, neuroinflammation, synaptic dysfunction, and synapse loss of AD and related dementia (*82, 83*). Building upon these findings, we reasoned that these data further provide a unique opportunity to identify human-specific regulatory elements responsible for maintaining the integrity of human cortical neurons and driving cortical neurogenesis.

We further explored regulatory elements driving cortical neurogenesis unique to humans using a previously compiled gene list that showed increased activity specifically in the developing human brain, when comparing gene expression between mice, macaques, and humans (*84*). Through overlapping human-gained enhancer (HGE) with snATAC-seq peaks from PiD and AD (Table S5), we identified an enhancer element that is both a differentially accessible peak in PiD and an HGE. Using chromatin co-accessibility analyses, we bioinformatically linked this differential accessible enhancer to *UBE3A*, even though it is located more than 40kbp away from its UTRs and around 80kbp away from its coding region (Fig. S8A). As a gene implicated in neuronal activity, *UBE3A* codes for a protein that plays a critical role in neuronal functioning, regulating proliferation and apoptosis (*85*). *UBE3A* loss-of-function mutation has been observed in individuals with Angelman Syndrome, while autism-linked *UBE3A* gain of function mutation was recently reported in a mouse model showing neurobehavioral deficits (*86, 87*). The cis-regulatory identified distal enhancers and HGE of *UBE3A* in neurons are more accessible in PiD (FDR-adjusted p-value = 4.40 ×10^−5^, Table S2C) (Figs. 2E, and 6A).

**Fig. 6.**
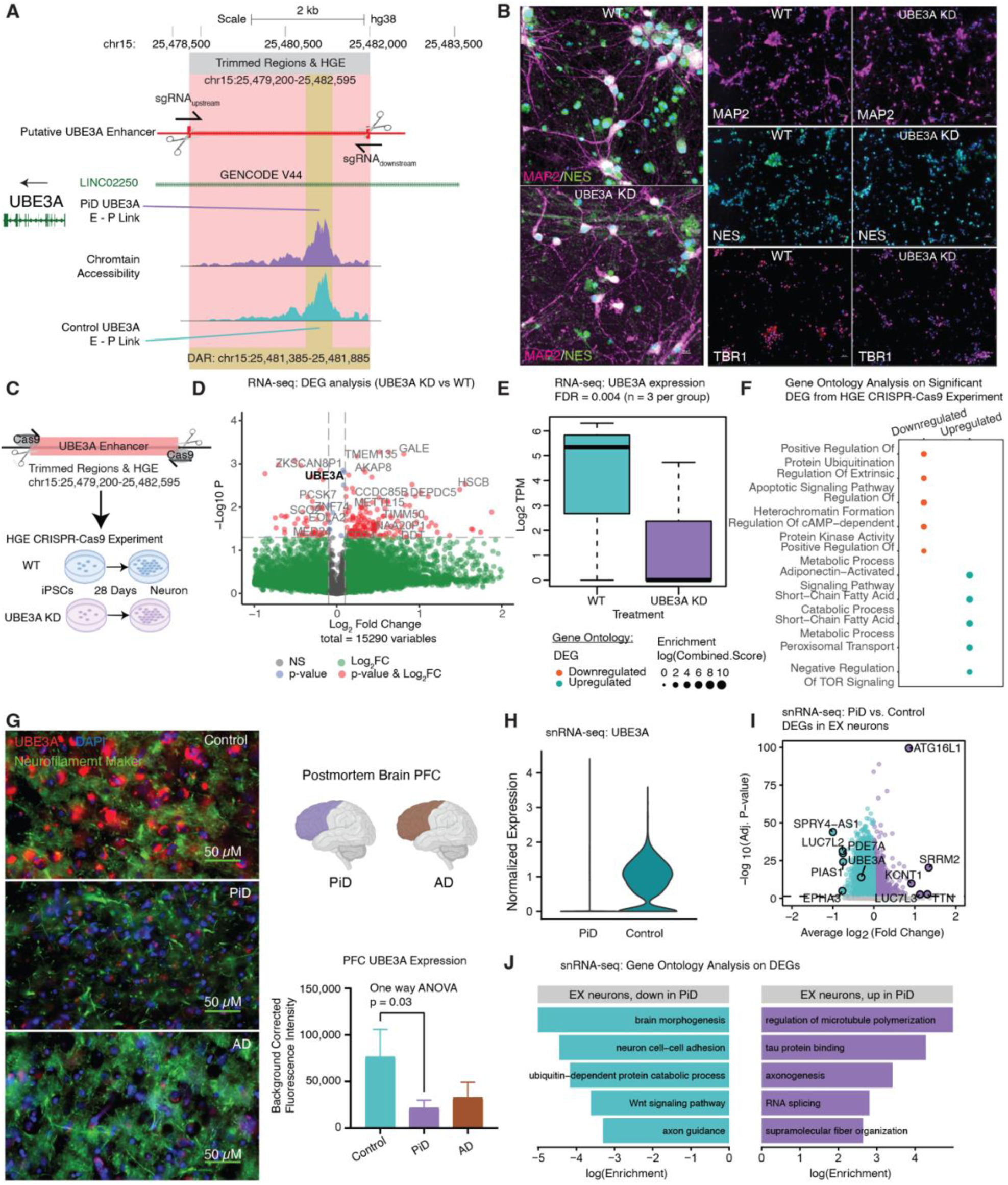
Mapping distal candidate cis-Regulatory Elements (cCREs) involved in synaptic function to their target genes. (**A**) Delta co-accessibility of UBE3A and enlarged CRISPR-edited enhancer regions of UBE3A in salmon and differentially accessible peaks in yellow overlap with intronic regions of long intergenic non-protein coding RNA 22 (LINC02250). (**B**) iPSC-derived neurons assessment, Left: Representative 40X images of 28-day WT and UBE3A KD cultures showing MAP2 (magenta) and NESTIN (green) expression. Scale bar = 10 µm. Right: Representative 20X images showing MAP2 (magenta), NESTIN (green), TBR1 (magenta), and Hoechst (blue) expression. Scale bar = 30 µm. (**C**) Experimental design for human-gained enhancer (HGE) CRISPR-Cas9 for UBE3A, RNA-seq performed on iPSC-derived neurons after 28 days of development. (**D**) Volcano plot of DEGs from RNA-seq (UBE3AKD vs WT), n = 3 per group. (**E**) UBE3A expression from RNA-seq (UBE3AKD vs WT). (**F**) Gene ontology of upregulated and downregulated DEGs from RNA-seq (UBE3AKD vs WT). (**G**) Representative immunofluorescence images for UBE3A (red), neurofilament marker (green), and DAPI (blue) from postmortem human brain tissue (PFC) of control (n=3), AD (n=5) and PiD (n=5) cases. 60X Images were captured using Nikon ECLIPSE Ti2 inverted microscope. Scale bar = 30 µm. (**H**) UBE3A expression from snRNA-seq in EX. (**I**) snRNA-seq DEG analyses in EX. (**J**) Gene ontology of upregulated and downregulated DEGs from snRNA-seq (PiD vs Control).

We hypothesized that the active HGE would enhance the expression of *UBE3A* or mitigate suppressive effects leading to its downregulation. Conversely, the elimination of this active HGE would presumably result in reduced levels of *UBE3A*. To validate whether this imputed enhancer is indeed the putative enhancer of *UBE3A*, we conducted CRISPR-edited experiments in iPSCs, wherein we targeted and excised the HGE region (chr15:25,479,200-25,482,595) (Fig. 6 A to C). CRISPR-modified (*UBE3A* KD) and isogenic control (WT) iPSCs were differentiated into cortical neurons using a modified *NGN2* induction protocol (*88*) (Methods). After 28 days, cortical neuron populations from both WT and *UBE3A* KD lines retained some *NES*-positive neural progenitor cells (Fig. 6B), and roughly 75% of nuclei co-localized with the mature neuronal marker *MAP2* with no statistically significant difference noted with *UBE3A* KD (p-value = 0.5687, unpaired t-test, two-tailed; Fig. S8B). Additionally, there was no notable expression difference in the early cortical layer marker, *TBR1*, seen with *UBE3A* KD (p-value = 0.8135, unpaired t-test, two-tailed; Fig. S8C), at greater than 50% in both populations. No substantial differences in marker expression were observed, confirming the neuronal identity of both the edited and unedited lines. In theory, if the predicted enhancer does regulate gene activity, removing it would interfere with its control mechanisms, resulting in reduced activity of the target gene. This approach has previously been employed to identify enhancers that regulate neocortical development (*89*). RNA-seq analyses performed on 28 days in vitro neurons revealed downregulation of *UBE3A* in the *UBE3A* KD neurons, confirming the predicted *UBE3A* HGE region regulates *UBE3A* expression (Fig. 6 D and E), and that the perturbation of *UBE3A* expression affected the expression of other genes (Fig. 6D). These genes are associated with the downregulation of protein ubiquitination, apoptosis, heterochromatin organization, cAMP-dependent protein kinase activity, and disruptions in various metabolic processes (Fig. 6F).

Given the intricate nature of human tissue, particularly in the context of disease conditions, our subsequent analyses in data derived from human tissue noted an enriched activity of chromatin accessibility (average log2FC > 0, Table S2C) for all distal peaks associated with *UBE3A* in the EX. Despite this, we observed a decrease in the proteomic and transcriptomic levels of *UBE3A*. In our immunofluorescence staining of *UBE3A*, we noted a statistically significant decrease in *UBE3A* levels in human PFC (Fig. 6G). Furthermore, our analyses of snRNA-seq DEGs in PiD also revealed *UBE3A* as one of the downregulated genes (Fig. 6 H and I). In our gene ontology analyses, we found that the downregulated genes were involved in various processes related to neuronal integrity, brain morphogenesis, neuron cell-cell adhesion, axon guidance, cell fate determination via the *Wnt* signaling pathway, and *UBE3A*-related ubiquitin-dependent protein catabolic processes. Conversely, among the upregulated terms, we observed enrichment in processes related to microtubule organization and tau protein regulation (Fig. 6J).

The discordance between increased snATAC-seq enhancer signal and decreased snRNA-seq gene expression for *UBE3A* may be attributed to a regulatory phenomenon where the chromatin region becomes more accessible to counteract the downregulation of its target genes. At the subcluster level, our integrated analysis of EX revealed consistent downregulation of *UBE3A* gene expression accompanied by increased chromatin accessibility at the *UBE3A* enhancer across most subclusters (Fig. S9 A to C). Subclusters EX4 and EX8, where fewer than 5% of cells exhibited accessible *UBE3A* enhancers, were excluded from this interpretation. The remaining subclusters (EX1 to EX3, EX5 to EX7) showed a consistent pattern of decreased UBE3A expression alongside elevated enhancer accessibility. This observation suggests that the discrepancy between increased snATAC-seq enhancer signal and decreased snRNA-seq gene expression for *UBE3A* cannot be explained solely by subcluster-specific differences, indicating the involvement of broader regulatory mechanisms. These findings underscore the complexity of regulatory dynamics within the disease context and highlight the need for further investigation into the regulatory processes underlying these observations.

## Discussion

Single-cell sequencing has been used to characterize the cell-type and cell state-specific changes in Alzheimer’s disease pathology extensively. While recent efforts have extended these approaches to other tauopathies (*7, 90, 91*), they remain comparatively understudied, particularly in Pick’s disease. In this study, we generated single-nucleus epigenomic and transcriptomic data from postmortem human brain tissue samples of Pick’s disease and cognitively normal controls. By integrating the analyses on cis- and trans-regulatory mechanisms with gene expression data, our approach at single-cell resolution enabled us to investigate the cellular diversity of the human PFC to compare shared and distinct regulatory mechanisms between these two tauopathies in excitatory neurons, astrocytes, microglia, and oligodendrocytes, and pinpoint the cell-type specific, disease-associated alterations. Meta-analyses in genome-wide association studies, supplemented with the assistance of snATAC and snRNA data, utilized AD and FTD GWAS genes and revealed putative and dysregulated risk genes for PiD. Systematic analyses of alteration in TF binding activity on promoter-enhancer links in both a genome-wide scale and gene-local region in PiD and AD revealed distinct and shared TF-regulatory networks from neurons and glial cells. Our single-nucleus data and customized approach to investigating cis- and trans-regulatory mechanisms altered in PiD and AD pathology led to the creation of an online interactive database, scROAD, which researchers are free to explore. We additionally generated RNA-seq data from iPSC-derived neurons following CRISPR-Cas9 editing, allowing us to validate imputed promoter-enhancer regulatory linkage from possible target genes involved in disease progression.

Although the precise molecular mechanisms driving PiD pathology remain elusive, our study provides insights into the intricate landscape of gene regulation in PiD, particularly the challenges in interpreting distal regulatory elements. Our differential analyses highlight the utility of our identified promoter-enhancer links in elucidating regulatory mechanisms and revealed widespread chromatin accessibility and gene expression changes linked to PiD and AD pathology across major cell-types. These alterations, spanning chromatin accessibility and expression of genes tied to synaptic signaling, apoptotic pathways, neuronal activity regulation, cellular stress responses, and intercellular communication, may indicate compensatory neuron-oligodendrocyte crosstalk that attempts to re-establish homeostasis by differentially modulating specific gene programs. Some promoter-enhancer connections facilitated increased chromatin accessibility, potentially serving as a compensatory mechanism to mitigate the dysregulation of target genes. Other alterations, including positive regulation of endocytosis, genes responsible for cellular metabolic processes, and genes encoding cellular response to unfolded/misfolded protein in astrocytes and microglia, may contribute to glial cell differentiation or immune activation in PiD and AD. Disruptions in the metabolic processes and cellular stress response compromise the balance in the cellular microenvironment and consequently contribute to the progression of PiD and AD.

While the causative molecular mechanisms of PiD remain unknown, our work offers insights that assist in unraveling the nature of gene regulation in PiD, especially regarding genomic loci with well-described heritable disease risk. We capitalized on the AD and FTD GWAS data to identify genes associated with phenotypic variability between PiD and AD because of similar pathological and clinical traits, such as tauopathies and cognitive decline. GWAS have been widely used to enhance our understanding of polygenic human traits and to reveal clinically relevant risk variants for neurodegeneration. Notably, we identified genetic risk variants that overlapped with specific cell-types to narrow down the potential non-coding variants underlying disease susceptibility. Furthermore, our analysis revealed that AD GWAS genes showed a substantial overlap with differentially expressed genes in PiD cases, suggesting that these associations are not random. This highlights the potential convergent regulatory mechanisms that may be shared between PiD and AD, despite the distinct clinical manifestations. Although this method has enabled the investigation of cell-type-specific disease-associated regulatory mechanisms, key limitations of the snATAC-seq assay without variant calling in PiD samples leave the opportunity for future studies and improvements.

Cell demise constitutes a defining characteristic of neurodegenerative ailments, including Pick’s and Alzheimer’s disease. More pronounced alterations in chromatin accessibility and gene expression were observed in excitatory neurons and oligodendrocytes in PiD compared to AD. In agreement with a previously observed association of rapid progression and early disease onset in PiD compared to AD (*3, 33*), as well as spatiotemporal differences (*7*), we found an elevation in the fold change in chromatin accessibility of dysregulation among genes and TFs, especially in excitatory neurons. Additionally, in excitatory neurons from PiD, we observed a complex regulatory mechanism that downregulated genes strongly associated with increased chromatin-accessible regions for the same genes through cis-regulated promoter-enhancer links, including genes responsible for neuronal activity and signaling, for example, *UBE3A*. A major contribution of our study lies in the identification of cell-type-specific enhancer-promoter pairs, potentially facilitating gene-regulatory alterations in PiD and AD, along with the TFs likely to bind to these regulatory elements within the respective cell-types. Our investigation into cis-regulatory elements and DNA-binding motifs, particularly in regions harboring fine-mapped SNPs, has uncovered potential disruptions in regulatory relationships, exemplified by the anti-amyloidogenic gene *ADAM10*. These disruptions, proximal to disease-associated SNPs, may lead to diminished TF binding activity and subsequent dysregulation of target gene expression. Furthermore, our analyses utilized the gene-specific-enhancer-binding TFs’ information to construct a TF regulatory network in neurons and demonstrated alterations in PiD and AD. We also provide insights into the regulatory landscape of TFs in glial cells across PiD and AD. We identified differential binding activities of TFs, such as *SPI1*, known as a major AD GWAS risk gene in microglia and associated with its development, *JUND* in astrocytes, known for its strong correlations with pTau and amyloid-beta, and transcriptional suppressors *HES1* and *ZBTB33* in oligodendrocytes, shedding light on their potential roles in disease pathogenesis. Moreover, the downstream dysregulation of TFs and genes associated with the highlighted TFs, including *CX3CR1*, *MAPT*, and *FOXO1*, emphasizes the intricate regulatory mechanisms implicated in neurodegenerative processes, with some alterations shared between PiD and AD, while others are uniquely observed in either condition.

The identification of functional regulatory elements in human excitatory neurons and the validation of their functions in iPSC-derived neurons enhance our understanding of epigenomic discovery. Leveraging these findings, we identified human-specific regulatory elements crucial for maintaining the integrity of cortical neurons in a neurodegenerative disorder, providing valuable annotations. Subsequent CRISPR-edited experiments in iPSCs confirmed the regulatory role of a putative enhancer in *UBE3A* expression. Furthermore, our observation of enriched chromatin accessibility near *UBE3A* in excitatory neurons, despite decreased *UBE3A* expression in snRNA-seq, highlights the complexity of gene regulation in the context of disease.

This study represents an important step in identifying PiD risk genes and leveraging transcription factor occupancy to predict regulatory mechanisms but has notable limitations. Small sample sizes, especially for rare cell-types, limited the power of analyses, making it difficult to detect subtle gene expression changes and chromatin accessibility patterns at refined subcluster levels. Statistical noise and variability inherent to snRNA-seq and snATAC-seq data further complicated these analyses. DARs in snATAC-seq were not interpreted in isolation in our study; instead, they were used in conjunction with transcription factor (TF) differential binding analyses to support the identification of putative CREs linked to differentially expressed genes. This integrative approach mitigates the risk of overinterpreting noisy DARs and strengthens the biological relevance of our regulatory inferences. Nonetheless, to further improve statistical power and resolution, larger datasets and higher-resolution techniques will be essential to improve robustness and resolution in future studies. Discrepancies in library preparation between PiD and AD datasets could introduce biases in data interpretation. To mitigate this, we compared disease versus control data within each study, applied stringent quality control, and used normalization and batch effect correction to harmonize data while preserving biological signals. Additionally, the relatively high abundance of oligodendrocytes in our dataset may partially reflect technical factors related to nuclei isolation and capture efficiency; nonetheless, emerging evidence suggests that oligodendrocytes contribute to neurodegenerative processes, and future studies focusing on this cell-type may offer valuable insights into the pathogenesis of PiD and AD.

Survival bias is another key limitation. Although we calculated a neuronal vulnerability module score, per Mathys et al. (*60*), to account for surviving neurons, this issue remains a challenge in single-cell studies. While our analysis included both surviving and vulnerable neurons in the PiD dataset, further investigation in dedicated studies is needed. The absence of PiD-specific GWAS data presents another constraint, limiting the direct applicability of FTD GWAS fine-mapping results to PiD. Despite incorporating AD and FTD GWAS data for overlap analysis, PiD’s rarity and unique pathology underscore the need for targeted genetic studies. Technical limitations, such as insufficient sequencing coverage and challenges with PCR-based library preparation, also restricted our ability to analyze *MAPT* haplotypes (*92*), hindering a full exploration of the *MAPT* locus in PiD pathology.

Although the study highlights key regulatory dynamics, such as increased *UBE3A* enhancer accessibility, these changes do not always result in corresponding gene expression increases. Additional layers, such as nonsense-mediated mRNA decay (NMD) or disease-related chromatin changes like relaxation and heterochromatin loss, could intervene and complicate these relationships. Future studies integrating advanced multi-omic approaches, including chromatin conformation assays and proteomics, will be crucial for unraveling the complex interplay between chromatin accessibility, gene expression, and disease-associated regulatory mechanisms in PiD.

Overall, our findings offer critical insights into the regulatory landscapes of PiD and AD, underscoring the value of integrated genomic approaches in unraveling the molecular mechanisms underlying neurodegenerative disorders. By highlighting the intricate interplay between transcriptional regulation and disease progression, this work emphasizes the need for a deeper understanding of these regulatory networks as a foundation for developing targeted and effective therapeutic strategies.

## Materials and Methods

### Postmortem human brain tissue

Human postmortem frontal cortex brain samples were obtained from UCI MIND’s Alzheimer’s Disease Research Center (ADRC), Harvard and Mt. Sinai tissue repositories. All participants, or participants’ legal representatives, provided written informed consent for the study. 50 mg of tissue from each sample (n = 9 control brain and n = 7 Pick’s brain) was dissected and aliquoted into a 1.5 ml tube inside a prechilled tissue dissection box as described previously (*93*). Samples were also selected based upon several covariates, including age, sex, postmortem interval (PMI), and disease comorbidity. Sample information is available in Table S1.

### Immunofluorescence

PFA-fixed human postmortem brain tissues (PFC region) were sectioned at 30 µm using a cryotome (Leica SM2010R). Sections were then rehydrated and washed in 1X sterile PBS and permeabilized using 1X sodium citrate buffer pH 6.0 (heated at 95°C for 10 mins). After blocking with 3% BSA solution or serum, sections were incubated with diluted primary antibodies (as per manufacture’s recommendation) at 4°C overnight (IBA1 antibody; Cat #NC9288364; 1:1000; Fisher Scientific, GFAP Polyclonal Antibody; Cat #PA3-16727; 1:500; ThermoFisher, p-tau (AT8) Cat #MN1020; 1:250; ThermoFisher; UBE3A; Cat #10344-1-AP; 1:1000; Proteintech, Anti neurofilament protein; Cat #837904; 1:1000; Biolegend). Secondary antibodies were selected and diluted according to the manufacturer’s instruction and incubated for 1.5-2 hrs. Sections were then washed (3X with PBS), mounted and cover slipped using anti-fade mounting media. Slides were imaged (20x/40x/60x) using Nikon ECLIPSE Ti2 inverted microscope. Images from 3 randomly selected areas of each slice were used for analyses.

### snATAC-seq tissue processing and nuclei isolation

Frozen brain tissue pieces were placed in 500 µL chilled 0.1X Lysis Buffer (1X lysis buffer diluted with lysis dilution buffer; please refer to snATAC-seq protocol (*93*) for more details) and immediately homogenized 15 times using a pellet pestle (Fisherbrand™ Pellet Pestle™ Cordless Motor with RNase-Free Disposable Pellet Pestles, Cat#12-141-364). The homogenized tissues were then incubated for 15 mins followed by addition of 500 µl of chilled Wash Buffer and filtration through a 70 µm Cell Strainer (Miltenyi Biotech). In the next step, a sucrose gradient (Nuclei PURE Prep Nuclei Isolation Kit, Cat #NUC201-1KT, Sigma) was prepared and nuclei were spun at 13,000 x g for 45 minutes at 4°C. After centrifugation, the debris and myelin from the top of the sucrose gradient were removed. Nuclei were resuspended, washed, filtered (through a 40 µm cell strainer), counted (using a cell counter), and then incubated in a Transposition Mix.

### snATAC-seq library preparation and sequencing

Transposed nuclei were loaded on 10X Genomics Next GEM Chip H (10x Genomics) to generate single-cell GEMS. GEMs were then transferred, incubated, and cleaned for further processing. Single nuclei ATAC-seq libraries were prepared using the Chromium Single Cell ATAC v2 (10x Genomics) reagents kit as per the manufacturer’s instructions. Library size distribution and average fragment length of each library were assessed with Agilent TapeStation High Sensitivity D5000 ScreenTapes and the concentrations were quantified using a Qubit Fluorometer. Libraries were sequenced on a NovaSeq 6000 (Illumina) in paired-end mode (read1N: 50 cycles, index i7: 8, index i5:16 cycles, read 2N:50 cycles) to generate approximately 500 M reads per sample.

### snRNA-seq library preparation and sequencing

45-50 mg of fresh frozen brain tissue (PFC) was homogenized in EZ Lysis buffer (Cat #NUC101-1KT, Sigma-Aldrich) and incubated for 10 min on ice before being passed through a 70 µm filter. The fresh tube with filtered homogenate was then centrifuged at 500 g for 5 min at 4°C and resuspended in an additional 1 mL of lysis buffer. After another centrifugation samples were incubated in Nuclei Wash and Resuspension buffer (1xPBS, 1% BSA, 0.2U/l RNase inhibitor) for 5 min. To remove myelin contaminants and debris, we prepared sucrose gradients and centrifuged the tubes at 13,000 g for 45 min at 4°C. Next, a debris removal solution (Cat #130-109-398, Miltenyi Biotec) was added to the nuclei suspension (and centrifuged at 3,000g 10 mins at 4°C) for a second round of cleanup. Debris-free clean nuclei suspension was then diluted in nuclei buffer (with BSA and RNase) before processing with the Nuclei Fixation Kit (Parse Biosciences). After fixation and permeabilization, nuclei were cryopreserved with DMSO until the day of library preparation. Libraries were prepared using EVERCODE™ WT V3 kit (Parse Bioscience) and quantified using Qubit dsDNA HS assay kit (Cat #Q32851, Invitrogen). D5000 HS kit (Cat #5067-5592, Cat #5067-5593; Agilent) was used for measuring the average fragment length of each library. Libraries were sequenced using Illumina Novaseq 6000 S4 platform (paired-end sequencing) for a sequencing depth of 50,000 read pairs/nuclei.

### Human iPSCs

The ADRC76 iPSC line (*94*) was provided by the UCI ADRC Induced Pluripotent Stem Cell Core. ADRC76 was generated from fibroblasts from an 83-year-old, white, male with no known disease. CRISPR/Cas9 editing was performed by UCI’s Stem Cell Research Center CRISPR Core to generate a homozygous deletion of the UBE3A enhancer region. Two guide RNAs were designed to the UBE3A enhancer region (chr15:25,479,200-25,482,595) and delivered with Cas9 as a RNP complex via electroporation. Clone C-14 (UBE3A KD) was selected and used for all experiments. Sanger sequencing was used to confirm the deletion, revealing a 1bp allelic difference in the deletion due to NHEJ-based DNA repair. Karyotyping was performed by Cell Line Genetics to ensure genomic integrity after CRISPR/Cas9 editing. Immunocytochemistry was used to confirm expression of pluripotency markers OCT4, SOX2, and SSEA4.

### Cortical neuron pellet generation

Cortical neurons were generated as previously described (*88*) with some modifications. Induced pluripotent cell lines were maintained in mTeSR Plus medium (Stem Cell Technologies Cat #100-0276) on GelTrex basement membrane (ThermoFisher Cat #A1413302) and passaged using ReLeSR (Stem Cell Technologies Cat #100-0484) at 80% confluence in the presence of CEPT (Chroman1-Tocris Cat #7163, Emricasan-Seleck Chemicals Cat #S7775, Polyamine supplement - Sigma Cat #P8483, Trans-ISRIB-R&D Systems-5284) (*95*). UBE3A mutant and parental lines were transfected via Nucleofection (LONZA Cat #VPH-5022) of the PB-TO-hNGN2 (Addgene Cat #172115*) plasmid and purified in the presence of 200 ng/mL Puromycin (Invivogen ant-pr-1) until the majority of cells showed plasmid expression as determined by BFP expression. Once a high BFP expression had been established, iPSCs dissociated to single cell with Accutase (ThermoFisher Cat #NC9464543) and seeded at 1 x 10^6^ cells per GelTrex coated 6 well in Induction media: Knockout DMEM/F12 (ThermoFisher); N2 supplement 100X (ThermoFisher); non-essential amino acids 100X (ThermoFisher), and supplemented with Doxycycline at a final concentration of 1µM (Sigma) and CEPT. The medium was changed every day. After 3 days, Uridine (U) and Fluorodeoxyuridine (FdU) were both added at 1mM (Sigma Cat #3750, Sigma Cat #0503). On day 4, the induced cells were passaged as single cells with Accutase and seeded at 2 x 10^6^ cells per Poly-D-Lysine coated 6 wells (Sigma Cat #P6407) in Cortical Neuron Culture Medium 1 (CM1): 1:1 Knockout DMEM/F12: BrainPhys neuronal medium without Phenol-Red (STEMCELL Technologies); B27 supplement, 50X (ThermoFisher); BDNF (10 µg/ml, STEMCELL Technolgies) in PBS containing 0.1% BSA (ThermoFisher); NT-3 (10 µg/ml, Preprotech) in PBS containing 0.1% BSA, GDNF (10 µg/ml, STEMCELL Technologies) in PBS containing 0.1% BSA; laminin final con. 1 µg/ml (ThermoFisher), Doxycycline (1 µM), U (1 µM), and FdU (1 µM). Cells were maintained an additional 24 days with half media changes every 3-4 days first with CM1 (day 7), then Cortical Neuron Culture Medium 2 (CM2) starting at day 10. CM2: BrainPhys neuronal medium without Phenol-Red (STEMCELL Technologies); B27 supplement, 50X (ThermoFisher); BDNF (10 µg/ml, STEMCELL Technologies) in PBS containing 0.1% BSA (ThermoFisher); NT-3 (10 µg/ml, Preprotech) in PBS containing 0.1% BSA, GDNF (10 µg/ml, STEMCELL Technologies) in PBS containing 0.1% BSA; laminin final con. 1 µg/ml (ThermoFisher), Doxycycline (1 µM), U (1 µM), and FdU (1 µM). Three successive passages of each cell line were differentiated in parallel with pellets collected and flash froze for RNAseq at D0, D4, and D28 along with PFA fixed coverslips.

### Cortical differentiation immunocytochemistry and image analysis

After 28 days in culture, cortical neuron populations were fixed with 4% paraformaldehyde (Fisher Scientific #50980487) for 10 minutes at room temperature, then washed three times with PBS (Corning #21030CV). Cells were permeabilized with 0.3% Triton-X (Sigma #T8787) in PBS for 10 minutes and then blocked with 2% goat serum (ThermoFisher #16210-064), 3% BSA (ThermoFisher #15260-037), 0.1% Triton-X, and 0.3M Glycine (Fisher #BP381-1) in PBS for 1 hour at room temperature and then incubated in primary antibody diluted in block, overnight at 4°C (anti-Nestin (1:1000) Millipore MAB5326, anti-MAP2 (1:1000; Synaptic Systems 188004), anti-TBR1 (1:250; Abcam ab31940). Primary antibody was removed, and cells washed three times with PBS and then incubated for 1 hour in secondary antibody diluted 1:1000 in block, in the dark at room temperature (Alexa Fluor Goat IgG (H+L) Secondary Antibody, ThermoFisher Scientific). Cells were washed with PBS for three times and then washed in PBS containing Hoechst 33342 (Sigma #14533) for 10 minutes and then a final wash in PBS. Coverslips were mounted with Fluoromount-G® (Fisher #OB10001) and allowed to dry. 40x images were acquired with an Olympus FLUOVIEW FV 3000 confocal microscope and 20x images were acquired at 20X on a Keyence BZ-X810 Widefield Microscope, 4 random images were taken per coverslip from each replicate differentiation. TBR1 positive cells and total nuclei (Hoechst) were quantified using Imaris Spots tool (Imaris Single Full software, BITPLANE) while MAP2 area was analyzed using Imaris Surface tool and the colocalization tool was used to count the number of nuclei were within the MAP2 positive surface. Both MAP2 positive nuclei and TBR1 positive cell counts were normalized by total nuclei per image. TBR1 and MAP2 expression values were analyzed using GraphPad Prism software using a Student’s two-tailed t-test, assuming equal variance.

### RNAseq experiments with iPSC neurons

Total RNA was extracted from iPSC-derived neurons using the Direct-zol RNA Miniprep kit (Zymo Research) following the manufacturer’s protocol. RNA quantity was measured using Qubit Fluorometric Quantitation, and RNA integrity was assessed using the RNA Integrity Number (RIN) on an Agilent 4100 Tapestation. Stranded Total RNA-Seq libraries were prepared using EvoPlus V2 kits (Roche), multiplexed, and sequenced on an Illumina platform to an average depth of approximately 50 million reads per sample. Raw FASTQ files were aligned to the human reference genome (GRCh38) using RNA-STAR (v 2.7), and transcript abundances were quantified in transcripts per million (TPM) using Salmon (v 1.10). Genes with TPM values greater than 1 in at least 20% of the samples were selected for downstream analysis. Differential gene expression analysis was performed using a linear regression model that accounted for batch effects, including those from library preparation and sequencing.

### Processing snATAC-seq data

We used Cellranger-atac count (v 2.0.0) to map raw snATAC-seq reads to the GRCh38 reference genome (downloaded from the 10x Genomics website) in each sample, quantifying chromatin accessibility for each cell barcode. First, we used the ArchR function createArrowFiles to format the output of Cellranger-atac, removing barcodes with transcription start site (TSS) enrichment less than 4 and fewer than 1000 fragments. This function also yields a barcodes-by-genomic-bins “tile matrix” and a “gene score matrix” which aggregates chromatin accessibility information proximal to each gene. We next used the R package ArchRtoSignac (*93*) to convert our dataset from ArchR to Signac format to proceed with downstream analyses in Signac. We next performed analyses of our recently generated snATAC-seq samples from PiD donors and cognitively normal controls with our previous snATAC-seq dataset of AD donors and controls as the reference dataset. We filtered snATAC-seq data with thresholds of TSS enrichment > 5 and fragment counts > 1500, similar to with the high-quality data standards established by Xiong et al (*27*). Following this, we created a merged object of the PiD and AD snATAC-seq datasets and generated an integrated, dimensionally-reduced representation using the Seurat function *FindIntegrationAnchors*, with reciprocal latent semantic indexing (RLSI) as the dimensionality reduction method. Using this anchor set, we performed transfer learning with the Seurat function *FindTransferAnchors* to predict cell-type identities for nuclei in the PiD dataset, based on annotations from the AD dataset. This transfer learning analysis assigned a probability score to each nucleus in the PiD dataset for its cell-type assignment. While some nuclei were confidently mapped to a single cell-type, others showed ambiguous mappings across multiple cell-types. To ensure high-confidence mappings, we filtered the PiD dataset to include only nuclei with a maximum prediction probability of 0.95. Subsequently, we conducted a final integrated analysis using LSI dimensionality reduction and Harmony, incorporating the biological sample as a covariate to account for batch effects. To ensure a fair and accurate comparison between PiD and AD, we implemented several measures, including rigorous batch correction and evaluation of UMAP visualizations of cell-type distributions across datasets. These steps confirmed that the observed contrasts between case and control groups were not confounded by batch effects, enabling robust comparative analyses of cell-type distributions. But to avoid potential confusion in our experimental design and comparison, we re-plotted UMAPs (Fig. S1C, Figs. 1, B to D) for PiD and AD separately against their respective control groups.

### Processing snRNA-seq data

We used split-pipe ParseBio pipeline (v 1.0.3) to map snRNA-seq reads to the GRCh38 reference transcriptome (downloaded from the Ensembl website) in each sample, quantifying unique molecular identifiers (UMI) for each cell barcode. Next, we accounted for potential ambient RNA contamination by applying Cellbender remove-background (v 0.2.0) to model the ambient signal and remove it from the UMI counts matrix for each sample. We then identified barcodes mapping to multiple nuclei (multiplets) by applying Scrublet (v 0.2.3) with default settings to each sample. We applied an initial quality control (QC) filter to remove barcodes with fewer than 250 UMI. Further, we applied sample-specific filters to remove barcodes in the top 5% of UMI, the percentage of mitochondrial reads, and the multiplet score within each sample. We finally applied a dataset-wide cutoff to remove barcodes with greater than 20,000 UMI, greater than 0.2 multiplet score, and greater than 5% mitochondrial reads, resulting in 68,999 barcodes for clustering analysis. We next performed clustering analysis with Scanpy with the following steps. First, we normalized gene expression for each cell by the total UMI counts in all genes and log transform using *sc.pp.normalize* total and sc.pp.log1p. Second, we performed feature selection using *sc.pp.highly_variable_genes* using the “Seurat v3” option for the feature selection method, retaining 3,000 genes for downstream analyses. Third, we scaled the normalized expression matrix for these 3,000 genes to unit variance and centered at zero mean using the *sc.pp.scale* function. Fourth, we performed linear dimensionality reduction with principal component analysis (PCA) using the *sc.tl.pca* function, which we then corrected on the basis of the sample of origin using Harmony. Fifth, we constructed a cell neighborhood graph using the top 30 harmonized PCs using *sc.pp.neighbors* function. We visualized this cell neighborhood graph using UMAP with the function *sc.tl.umap*. We performed an initial round of Leiden clustering with a high-resolution parameter (resolution = 3) to reveal additional clusters of low-quality cells which may have escaped our previous QC filtering, and to annotate major cell-types based on a panel of canonical marker genes. After removing two low-quality clusters, we split apart the dataset by major cell lineages (excitatory neurons, inhibitory neurons, oligodendrocytes, and astrocytes) to perform sub-clustering analyses, yielding our final clustered and processed snRNA-seq dataset.

### Differential accessible open chromatin analyses

We systematically performed the analyses of differential open chromatin accessibility across each cellular type. This involved contrasting the disease states with their respective control conditions. For all the differential analyses employed, differentially accessible peak scrutiny was facilitated by implementing logistic regression (*test.use = ‘LR’*) to draw comparisons between cellular groupings. Logistics regression was utilized based on the accessibility interface of a specified open chromatin region (OCR) within varying groups of the selected cell-type. This is a protocol recommended by the Signac package (v 1.9.0) (*96*). The differential analyses were executed in Signac by deploying the same *FindMarkers* function found in Seurat (v 4.3.0). The accessible peaks that exhibited an adjusted p-value (corrected by Bonferroni method) of less than 0.05, accompanied by a minimum cellular fraction (min.pct > 0.05) in either of the two groups, were categorized as differentially accessible peak between the cellular groupings. We ran a comparative analysis of chromatin accessibility between the two diagnosis groups, specifically Pick’s disease (PiD) and Alzheimer’s disease (AD), and their age-appropriate cognitively normal counterparts. This was conducted within the human single-nucleus ATAC-seq dataset. The differential accessibility findings were visualized using a Complexheatmap (*97*), divided by diagnosis comparison and hierarchically aggregated based on the avg log2FC of differentially accessible peaks. This enabled us to focus on changes specific to each cellular type within each genomic classification. Finally, to single out the biological pathways and processes exhibiting a notable enrichment within our promoter differentially accessible peak sets or promoters of cis-regulatory-associated differentially accessible peaks present in distal and intronic regions, we invoked the support of the GREAT R package (v 2.0.2) (*98, 99*). Source of variation analysis was conducted using the variancePartition R package (v 1.32.5) (*100*) to assess the contribution of experimental variables to variation in both gene expression and chromatin accessibility in single-cell data, and to inform covariate selection in both the differential gene expression and differential chromatin accessibility models (Fig. S10 A and B). In addition, we systematically tested the effect of including different covariates on differential chromatin accessibility outcomes to evaluate model sensitivity and potential overfitting (Fig. S10 C to G).

### Differential gene expression analyses

We identified unbiased marker genes in each of our snRNA-seq clusters by a one-versus-all differential gene expression test using the Seurat (v 4.3.0) function *FindAllMarkers* with MAST as our differential expression model. We used sequencing biological sample and total number of UMIs per cell as model covariates. We performed differential expression analyses to compare gene expression signatures in cells from PiD and control samples in each of our major cell-types (excitatory neurons, inhibitory neurons, oligodendrocytes, OPCs, astrocytes, pericytes, endothelial cells, and microglia). Similar to our cluster marker gene test, we used MAST as our differential expression module with biological sample, sex, and number of UMI as model covariates. We used the R package enrichR (v 3.0) to perform pathway enrichment analyses for the DEGs in our excitatory neuron population.

### Statistical fine-mapping of candidate causal variants residue within cell-type specific accessible peaks from the snATAC-seq data

We sourced comprehensive genome-wide association studies (GWAS) pertinent to Alzheimer’s Disease (AD) (*12*) and frontotemporal degeneration (FTD) (*13*). The summary data pertaining to the AD GWAS was procured from the European Bioinformatics Institute GWAS Catalog (accession number: GCST90027158), whilst the FTD GWAS summary data was retrieved from the International Frontotemporal Dementia Genetics Consortium. To streamline the output files of the GWAS summary statistics from each dataset, we employed a uniformly designed pipeline, MungeSumstats (*101*). The application of this tool was governed by parameters that have been specified comprehensively in our GitHub repository. To further elucidate the role of single nucleotide polymorphisms (SNPs) pertaining to AD, we fine-mapped these SNPs within a 1-Mb window of the lead variants of AD risk loci that had been unearthed in the initial GWAS investigation (*12*). In addition to the AD SNPs, the detection of lead SNPs associated with FTD (*13*) required the identification of specific genetic markers encased within a 1-Mb spectrum present on all chromosomes. The selection criteria for these markers were established based on the statistical significance of their corresponding p-values. To accommodate all SNPs within the linkage disequilibrium (LD) block, we estimated pairwise LD between SNPs within the 1-Mb window of the GWAS lead variant. This estimation was performed using PLINK (v 1.9 and v 2.0) (*102*). Once the lead SNPs from the FTD and AD GWAS had been secured, the identified data was customized according to the corresponding 1-MB range LD matrix, within the sparse multiple regression model. This model was then implemented in the fine-mapping instrument, Sum of Single Effects (SuSiE) (*103, 104*). We managed to acquire a number of credible sets (CSs) for identified FTD and AD GWAS risk loci with high probability (a posterior inclusion probability: PIP > 0.95). In order to prioritize these credible sets, we aligned SNP locations with our snATAC-seq open chromatin regions. The fine-mapped casual SNPs within the identified cell-types were assessed for credibility by cross-referencing the GWAS risk genes’ expression level across all cell-types using control data from published resources (*17, 20, 39*). The final step included checking two scores - the probability of being loss-of-function intolerant (pLI) and the loss-of-function observed/expected upper-bound fraction (LOEUF) for the prioritized GWAS risk loci. These scores reflect the integrity of a gene or transcript in tolerating protein truncating variation (*38*).

### Utilizing publicly available datasets

We obtained the sequence data from three peer-reviewed single-nucleus RNA sequencing (snRNA-seq) studies related to Alzheimer’s Disease (AD) (*17, 20, 39*). The datasets represented in the works of Mathys et al. (2019), and Zhou et al. (2020) were accessed via the Synapse platform (referenced under syn18485175 and syn21670836, respectively). In the context of the dataset for the Morabito et al. (2021) study, which was formulated by our research team, a download was not necessitated, but you can access it under syn22079621. Although the pipeline was largely consistent, there were minor deviations in terms of parameter adjustments as per the individual requirements of each dataset. A comprehensive delineation of these nuanced changes is documented in our GitHub repository. Additionally, to enhance the reliability of our analyses, we incorporated PsychENCODE scQTL datasets from Pratt et al. (*34*) and Emani et al. (*35*), DLPFC snRNA-seq scQTL data from Fujita et al. (*36*), and neuronal and glial caQTL data from Zeng et al. (*37*). These datasets were leveraged to validate key findings from our snATAC-seq analyses. Specifically, we focused on significant eQTLs (p-value < 1×10^-5) associated with fine-mapped GWAS genes within the TF regulatory networks identified in this study. Furthermore, we cross-referenced our peak sets with Xiong et al.’s epigenomic data (*27*), further substantiating the robustness of our findings.

### Finding co-accessible peaks with Cicero to establish putative enhancer-promoter linkage

We initiated the conversion of the SeuratObject into the CellDataSet framework utilizing the *as.cell_data_set* function offered within the SeuratWrappers toolkit (v 0.3.0). This was subsequently transformed into a Cicero object through the application of the *make_cicero_cds* function taken from the Cicero package (v 1.3.4.11). The *run_cicero* function, a key component of the Cicero suite, was then employed to calibrate the co-accessibility of open chromatin peaks across the genome for each cell-type. The predominant objective here was to predict cis-regulatory interactions within a genomic window of 300,000 base pairs. The construction of a linkage co-accessibility score for each associating pair of accessible peaks was completed using a graphical LASSO regression model. This package and approach were based on techniques detailed in the Cicero method (*31*). The understanding being that an increased co-accessibility score denoted a stronger bond between an OCR pair and hence, greater confidence could be assigned to this pairing within a given dataset. Within the total ensemble of OCR pairs, we prioritized our examination on pairs identified as enhancer-promoter. The rationale for this selective focus stemmed from the potential for the enhancer-enhancer pair’s co-accessibility score to originate from inherent enhancer-enhancer interactions. This in turn could lead to a perceivable reduction in the co-accessibility scoring for the enhancer-promoter pair. Lastly, a comparative study was undertaken to calculate the delta co-accessibility score within identical OCR pairs. In this step, diseased states were compared with their corresponding control settings. The purpose of this comparison was to highlight any enhanced enhancer-promoter linkages that could potentially be contributing to the advancement of the disease.

### Characterizing biological functions of putative enhancer-promoter linkage

We used NMF (v 0.23.0) as implemented in the R NMF package using k = 25 matrix factors on the cis-regulatory-linked-enhancer accessibility matrix averaged by each snATAC-seq cluster split by cells from PiD control and PiD samples, as well as AD control and AD samples, yielding 25 enhancer modules. The NMF basis matrix (W) was used to assign each enhancer to its top associated module, and the NMF coefficient matrix (H) was used to determine which cell cluster that each module was most associated with. We applied NMF with a factorization rank of k = 25 to decompose the enhancer-by-cluster matrix derived from cis-regulatory-linked enhancer accessibility values. While the factorization yielded 25 enhancer modules, Fig. 2B highlights a focused subset of 9 modules that exhibited clear cell-type-specific accessibility patterns most relevant to the aims of the figure. The remaining modules were not excluded from downstream analysis, but were omitted from this particular plot to maintain visual clarity and focus. To identify biological processes associated with these enhancer modules, we used the enrichR (v 3.0) package to query enriched GO terms for the set of target genes in each enhancer module in the GO Biological Processes 2021, GO Cellular Component 2021, GO Molecular Function 2021 databases, Human WikiPathway 2021 and Human KEGG 2021.

### Transcription factor Occupancy prediction on snATAC-seq chromatin accessibility

TOBIAS (*56*) stands as a robust, precise, and rapid footprinting framework, facilitating a comprehensive exploration of TF binding occupancy for numerous TFs concurrently on a genome-wide scale as well as at the gene local region. We want to use this ATAC-seq analysis toolkit to investigate the kinetics of transcription factor (TF) binding in PiD, AD and their distinctions compared to respective control conditions, and we turn to TOBIAS for its capabilities as the ATAC-seq TF footprinting analyses toolkit. Our initial steps involved the extraction and categorization of cell barcodes based on both cell-type and diagnosis. Subsequently, we compiled distinct .bam files for each condition, serving as the requisite input format for the TOBIAS *ATACorrect* step. This particular tool within TOBIAS corrects the inherent insertion bias of Tn5 transposition. Following this correction process, the central task in footprinting commenced with the identification of protein binding regions across the entire genome. Utilizing single-base pair cut site tracks generated by *ATACorrect*, TOBIAS *FootprintScores* was employed to compute a continuous footprinting score across these regions. This approach enhances the prediction of binding for transcription factors even with lower footprintability, characterized by weaker footprints. Subsequently, the footprints were plotted using the function *PlotAggregate* to visualize and compare the aggregated signals across the specified conditions. This step serves to provide a tangible representation of TF binding occupancy and facilitates comparative analyses of these changes under different diagnosis conditions in each cell-type.

### Transcription factor Regulatory Network Construction

To construct a comprehensive TF regulatory network, we integrated insights from Cicero (*31*) and TOBIAS (*56*). First, leveraging Cicero, we focused on the predicted cis-regulatory interactions within a 300,000 base pair window, and grouped them based on the genomic class around its target gene identified by accessible promoter peaks. By prioritizing the examination of enhancer-promoter pairs within the ensemble of OCR pairs, we discerned potential interactions crucial for regulatory difference. Simultaneously, using TOBIAS, we explored TF binding activity in the selected gene local region by applying ATAC-seq footprinting analyses to identify protein binding regions across the genome. Following that, we utilized knowledge of accessible peaks’ cis-regulatory activity and gene local region TF binding activity to construct a TF regulatory network for selected target genes using R package igraph (v 2.0.1.9005). This approach combines co-accessibility from Cicero and footprinting from TOBIAS, providing a nuanced perspective on the regulatory landscape. The resultant TF regulatory network offers a multifaceted depiction of the interplay between TFs, enhancers, and promoters, enhancing our ability to decipher the intricacies of gene regulation in the context of PiD and AD. In the network, top-selected genes were filtered to include only those with an adjusted p-value < 0.05 and expressed in at least 5% of cells (pct > 0.05). From this set, we further prioritized genes within the top 50% of co-access scores derived from cis-regulatory link calculations, with TF binding on the cis-regulatory elements confirmed by TOBIAS package. Next, we retained the top 10 or fewer genes with the largest absolute log fold changes for both upregulated and downregulated genes associated with the selected transcription factors (TFs) in the TF network.

## Acknowledgments

We thank the International FTD-Genetics Consortium (IFGC) for summary data. We thank the UCI MIND’s ADRC and NIH NeuroBioBank for providing tissue. We thank the UCI Stem Cell Center and CRISPR Core for providing iPSC stem cells for CRISPR validation. We thank the UCI Genomics Research and Technology Hub for providing their facilities and sequencing our single-nucleus RNA-seq and ATAC-seq libraries. This work utilized the infrastructure for high-performance and high-throughput computing, research data storage and analyses, and scientific software tool integration built, operated, and updated by the Research Cyberinfrastructure Center (RCIC) at the University of California, Irvine (UCI).

## Funding

Fundings for this work were provided by:

National Institute on Aging grant 1RF1AG071683 (V.S.),

National Institute on Aging grant U54 AG054349-06 (MODEL-AD),

National Institute on Aging grant 3U19AG068054-02S (V.S.),

National Institute of Neurological Disorders and Stroke grant P01NS084974-06A1 (V.S.),

National Institutes of Health R35NS116872 (L.T.),

Adelson Medical Research Foundation funds (V.S.),

National Institute on Aging predoctoral fellowship T32AG000096-38 (E.M.),

National Institute on Aging predoctoral fellowship 1F31AG076308-01 (S.M.)

## Author contributions

Conceptualization: Z.S., S.D., S.M., E.M., and V.S.

Methodology: Z.S., S.D., S.M., J.S., E.M., and V.S.

Investigation: Z.S., S.D., S.M., and V.S.

Visualization: Z.S., S.D., and S.M.

Funding acquisition: V.S.

Project administration: V.S.

Supervision: V.S.

Validation: Z.S., S.D., N.E., J.S., A.S and L.T.

Software: Z.S., S.M., and A.C.

Formal analysis: Z.S., S.D., S.M., J.S., E.M., S.S., N.R., and Z.C.

Data curation: Z.S., S.D., J.S., A.C. and E.M.

Resources: E.H. and L.T.

Writing—original draft: Z.S., S.D., S.M.

Writing—review and editing: Z.S., S.M., E.M., and V.S.

## Competing interests

Authors declare that they have no competing interests.

## Data and materials availability

All data needed to evaluate the conclusions in this paper are present in the paper and/or the Supplementary Materials. All raw and processed single-nucleus ATAC, single-nucleus RNA and bulk RNA sequencing data in this manuscript have been deposited into the National Center for Biotechnology Information Gene Expression Omnibus (GEO) database (GSE259298). Code for the data analyses used for this study is available on Zenodo (DOI: 10.5281/zenodo.14649027). Supplemental tables have been deposited on the research data repository, Dryad (doi.org/10.5061/dryad.h9w0vt4t9). Additionally, we provide an interactive database, scROAD, to facilitate exploration of single-cell transcription factor occupancy and regulatory networks. Researchers can access scROAD (http://swaruplab.bio.uci.edu/scROAD). Access is unrestricted for noncommercial research purposes. Please note that any intended commercial use of this resource will require appropriate material transfer agreements (MTAs). The ADRC76 iPSC line can be provided by the Induced Pluripotent Stem Cell Core at the University of California, Irvine, pending scientific review and a completed MTA. Requests for the ADRC76 iPSC line should be submitted to the UCI Office of Research.

## Supplementary Materials

### Supplementary Text

#### scROAD Interactive Database

We have developed scROAD (Single-cell Regulatory Occupancy Archive in Dementia), an interactive online resource designed to explore single-cell cCREs (candidate cis-regulatory elements) transcription factor occupancy data. This database provides comprehensive information derived from snATAC-seq analysis of human postmortem prefrontal cortex (PFC) tissue, with a specific focus on Alzheimer’s Disease (AD) and Pick’s Disease (PiD). scROAD enables researchers to perform exploratory searches for genes, transcription factors (TFs), and SNPs, as well as to visualize transcription factor regulatory networks. Additionally, users can download the data for their own research purposes.

Accessible at http://swaruplab.bio.uci.edu/scROAD, this resource is freely available for noncommercial research purposes. By providing high-resolution regulatory data, scROAD supports the broader scientific community in advancing our understanding of transcriptional regulation in dementia and fosters the generation of novel hypotheses and discoveries.

**Fig. S1.**
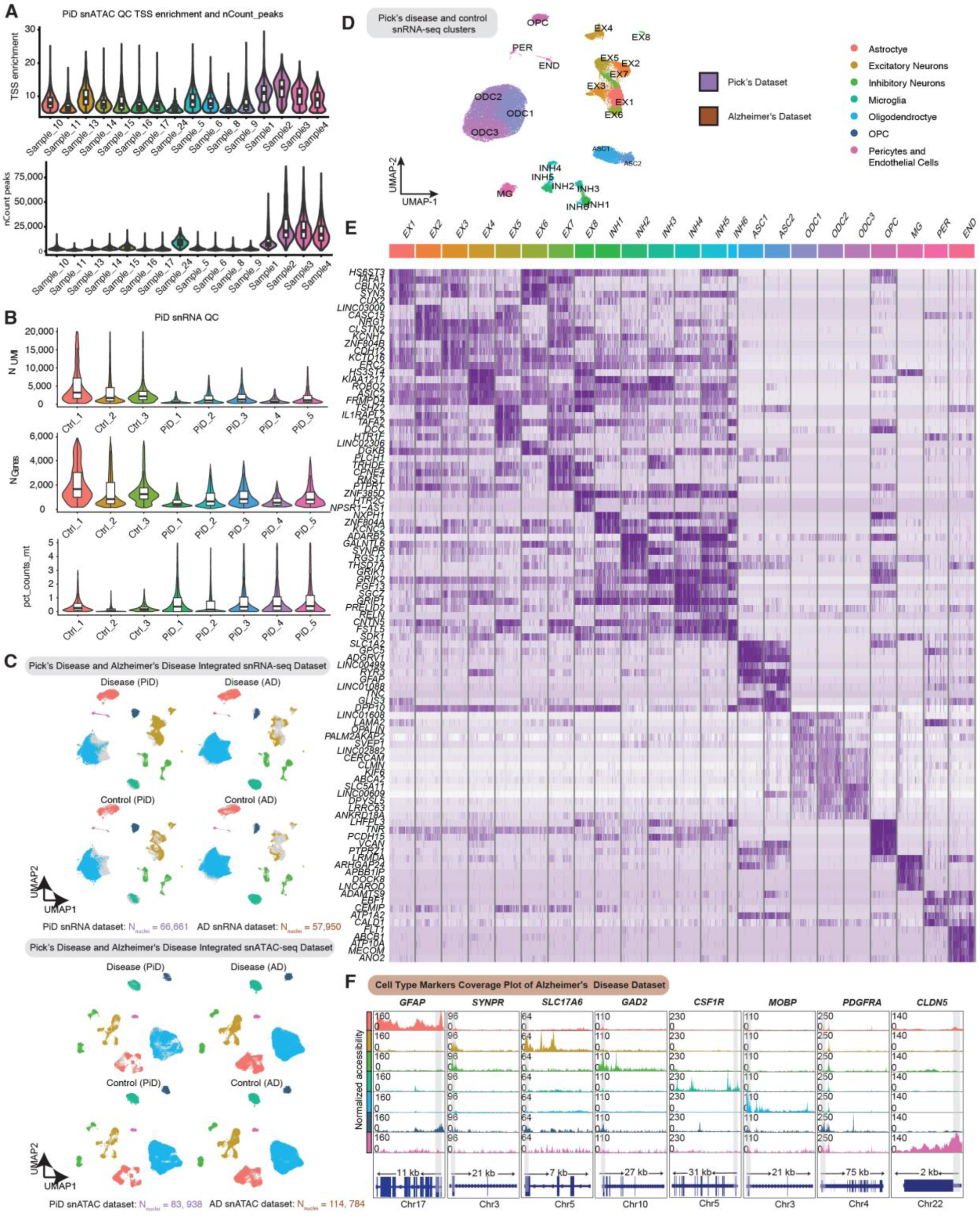
Quality control and cell type annotations of the PiD and AD snMulti-omic datasets. (**A**) Violin plot showing the number of peak counts in the samples from the PiD PFC snATAC-seq dataset. (**B**) Violin plot showing the number of UMI, genes and mitochondrial percentage in the samples from the PiD PFC snRNA-seq dataset. (**C**) Integrated Uniform Manifold Approximation and Projection (UMAP) visualizations by diagnosis for snRNA-seq and snATAC-seq data from PiD and AD. (**D**) Uniform Manifold Approximation and Projection (UMAP) visualizations for clusters of snRNA-seq data from PiD. (**E**) Heatmap of canonical cell-type markers for snRNA-seq data from PiD. (**F**) Coverage plots for canonical cell-type markers in AD dataset: *GFAP* (chr17:44905000-44916000) for astrocytes, *SYNPR* (chr3:63278010-63278510) for neurons, *SLC17A6* (chr11:22338004-22345067) for excitatory neurons, GAD2 (chr10:26214210-26241766) for inhibitory neurons, *CSF1R* (chr5:150056500-150087500) for microglia, *MOBP* (chr3:39467000-39488000) for oligodendrocytes, *PDGFRA* (chr4:54224871-54300000) for Pericytes and Endothelial cells in PiD dataset. The gray bar within each box highlights the promoter regions.

**Fig. S2.**
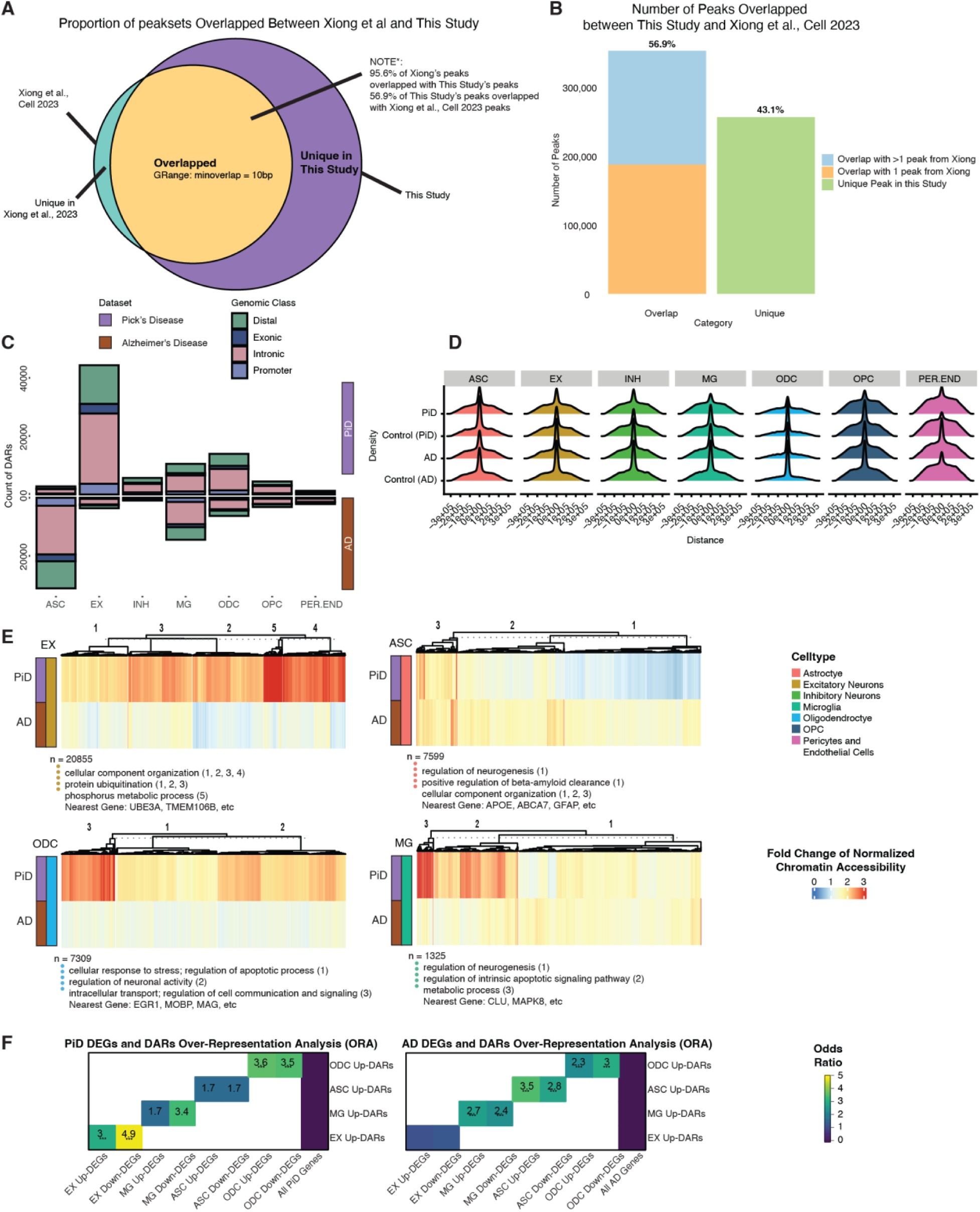
Comprehensive analysis of peak overlaps, cis-regulatory construction and changes in PiD and AD. (**A**) Proportion of peaksets overlapped between Xiong et al. (*27*) and this study. (**B**) Number of peaks overlapped between this study and Xiong et al. (**C**) Genomic type classification of differential open accessible regions grouped by cell types (P-value < 0.05) between PiD and AD with their respective controls. (**D**) Ridgeline plot showing the distance of imputed enhancers from the promoters. (**E**) Heatmaps of fold changes (Disease vs. Control) on normalized chromatin accessibility of differential accessible intronic regions in excitatory neurons, astrocytes, microglia and oligodendrocytes (FDR adjusted P-value < 0.05 and abs(log2FC) > 0.5), gene ontology acquired from GREAT and examples of promoters and distal regions’ cis-regulatory linked gene as in the panel of Fig. 2E. (**F**) Over-representation analysis (ORA) of DEGs (snRNA-seq) and DARs (snATAC-seq) from PiD and AD.

**Fig. S3.**
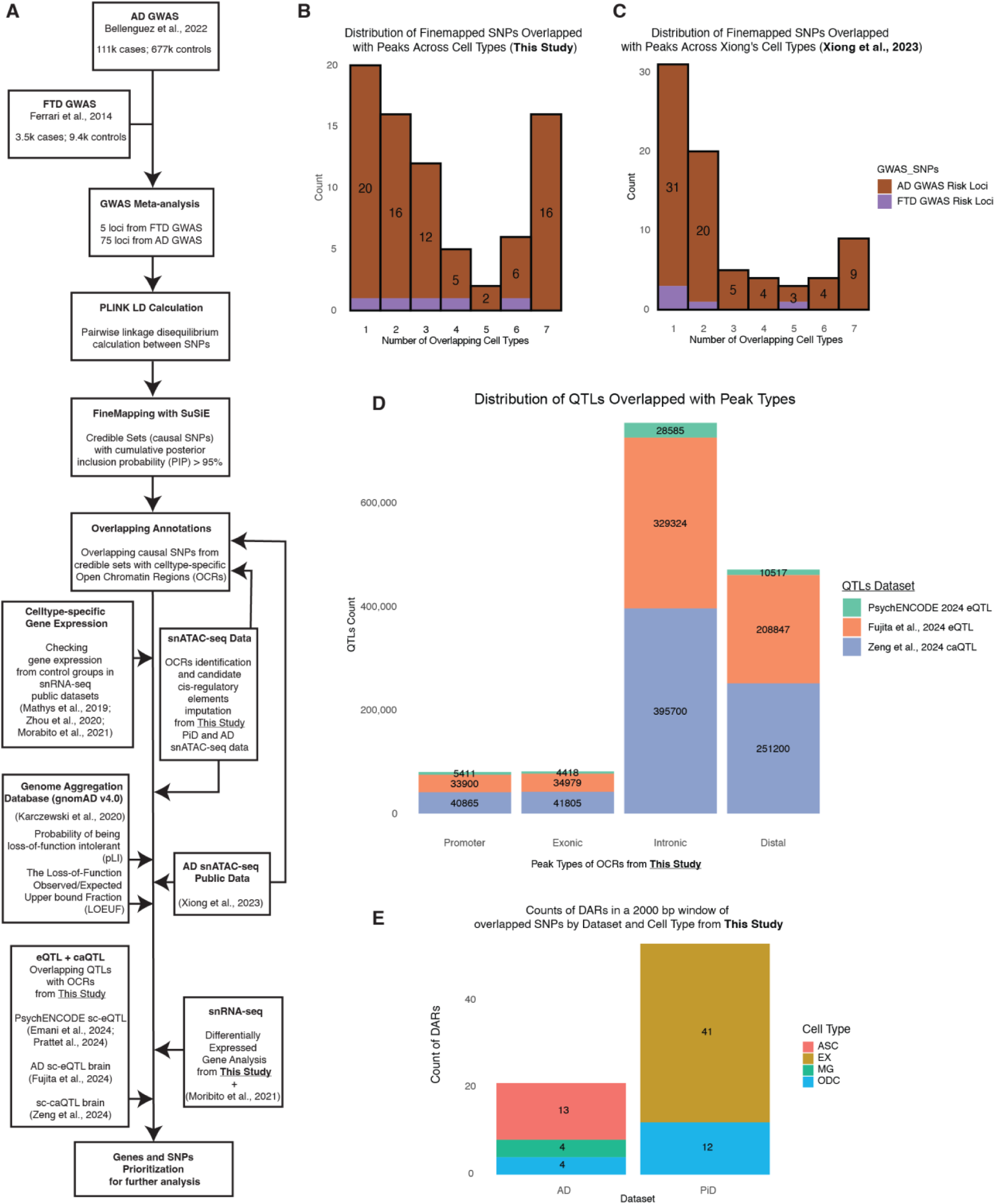
Integration and validation of GWAS fine-mapped SNPs with open chromatin regions in FTD and AD. (**A**) The schematic of analyses showing the summary of Frontotemporal Dementia (FTD) and Alzheimer’s Disease GWAS meta-analyses, fine-mapping, and other data processing steps to link causal SNPs to snATAC-seq accessible peaks in specific cell types. (**B**) Histogram of Overlapped Credible Sets with this study: Count vs. Overlapped Cell Types. It describes a histogram that displays the count of overlapped credible sets and their associated number of overlapped cell types. (**C**) Histogram of Overlapped Credible Sets with Xiong et al. (*27*): Count vs. Overlapped Cell Types. (**D**) Distribution of QTLs (*34–37*) overlapped with peak types from this study. (**E**) Counts of DARs in a 2000 bp window of overlapped SNPs by dataset and cell type from this study.

**Fig. S4.**
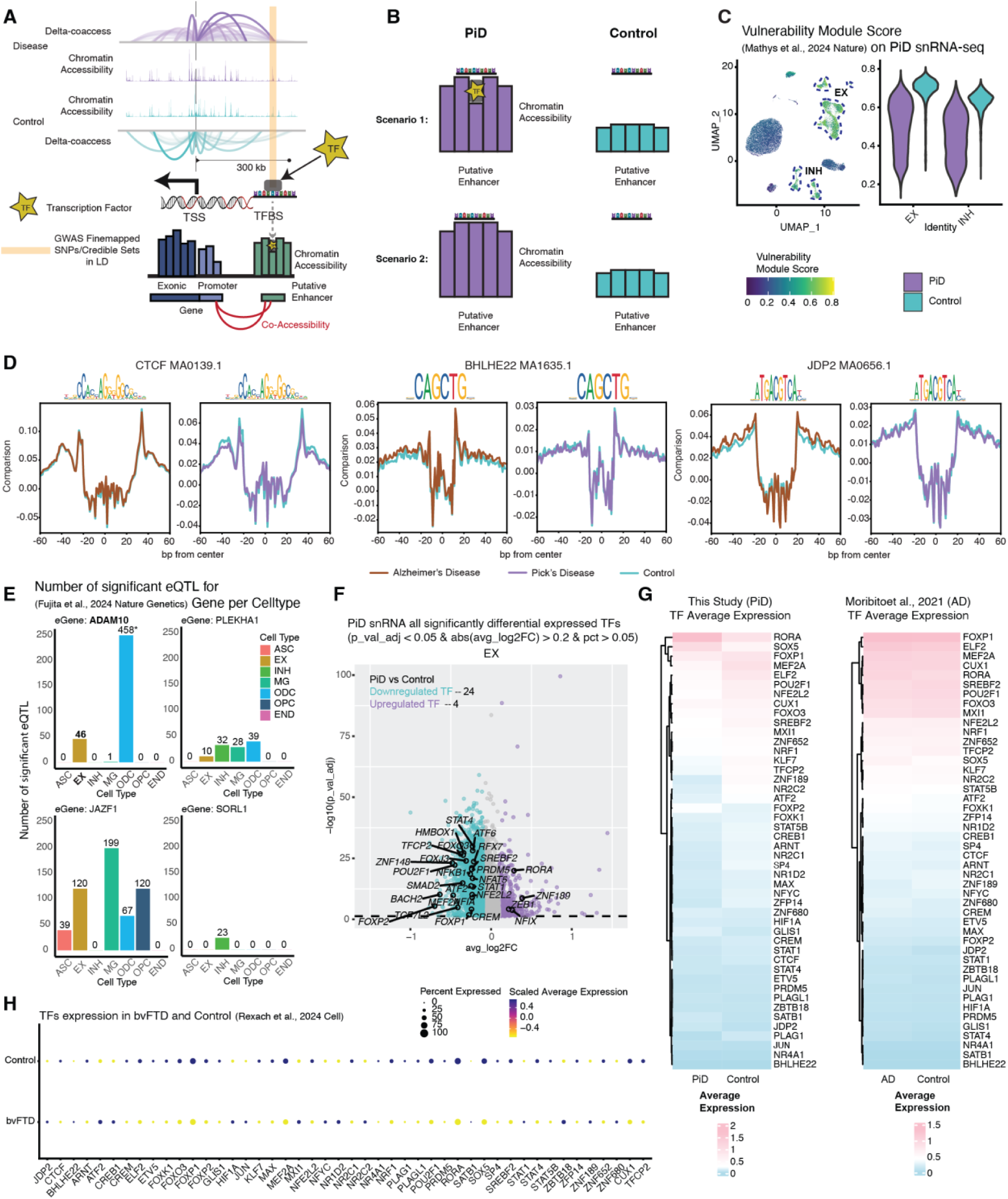
EX transcription factor dynamics and differential expression in PiD and AD. (**A**) Open chromatin co-accessibility plot with TF binding occupancy calculation. (**B**) Two open chromatin scenarios in PiD and control; scenario 1: Open chromatin with TF binding (footprint) in PiD; scenario 2: Open chromatin without TF binding (no footprint) in PiD. (**C**) Vulnerability module score (*60*) of neurons, EX and INH, in PiD. (**D**) Aggregated TF footprints of *CTCF* (MA0139.1), *BHLHE22* (MA1635.1), and *JDP2* (MA0656.1) in AD and PiD. (**E**) Significant AD sc-eQTLs (p-value < 1×10^-5) from Fujita et al. (*36*) by cell type. (**F**) PiD snRNA all significantly differential expressed TFs. (**G**) Heatmap of average expression difference of highlighted TFs and top selected differentially TFs regulated by highlighted TFs between PiD or AD with their matched control (FDR-adjusted p-value < 0.05). (**H**) Dotplot of differentially expressed TFs in Rexach et al. (*7*) bvFTD versus their respective controls.

**Fig. S5.**
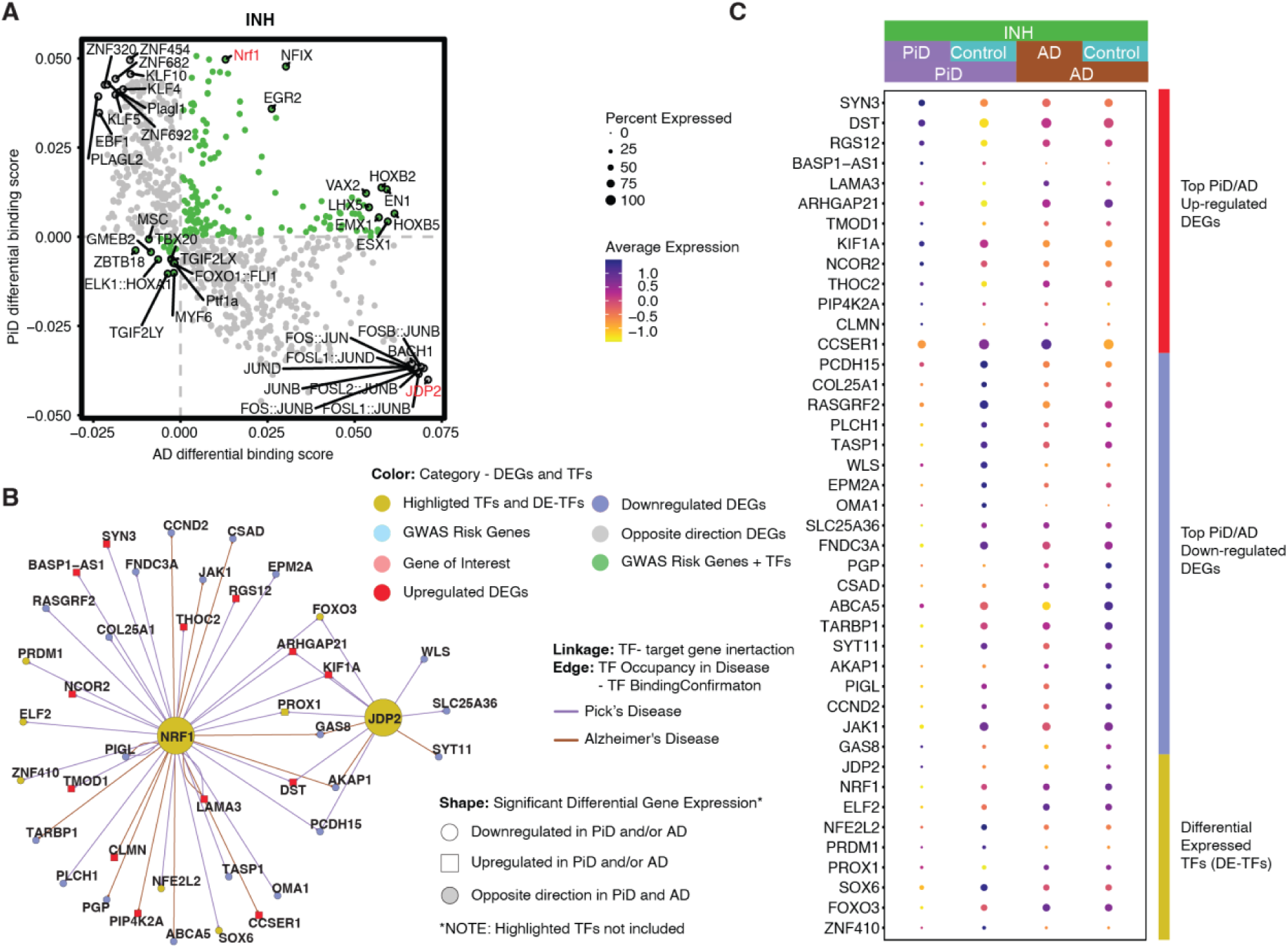
INH TF binding difference and regulatory networks in PiD and AD. (**A**) Genome-wide Tn5 bias-subtracted TF differential footprinting binding score of PiD and AD in INH. (**B**) *NRF1* and *JDP2* TF regulatory networks showing the predicted candidate target genes for INH. (**C**) Dot-plot of the differentially expressed genes and TFs in PiD and AD versus their respective controls.

**Fig. S6.**
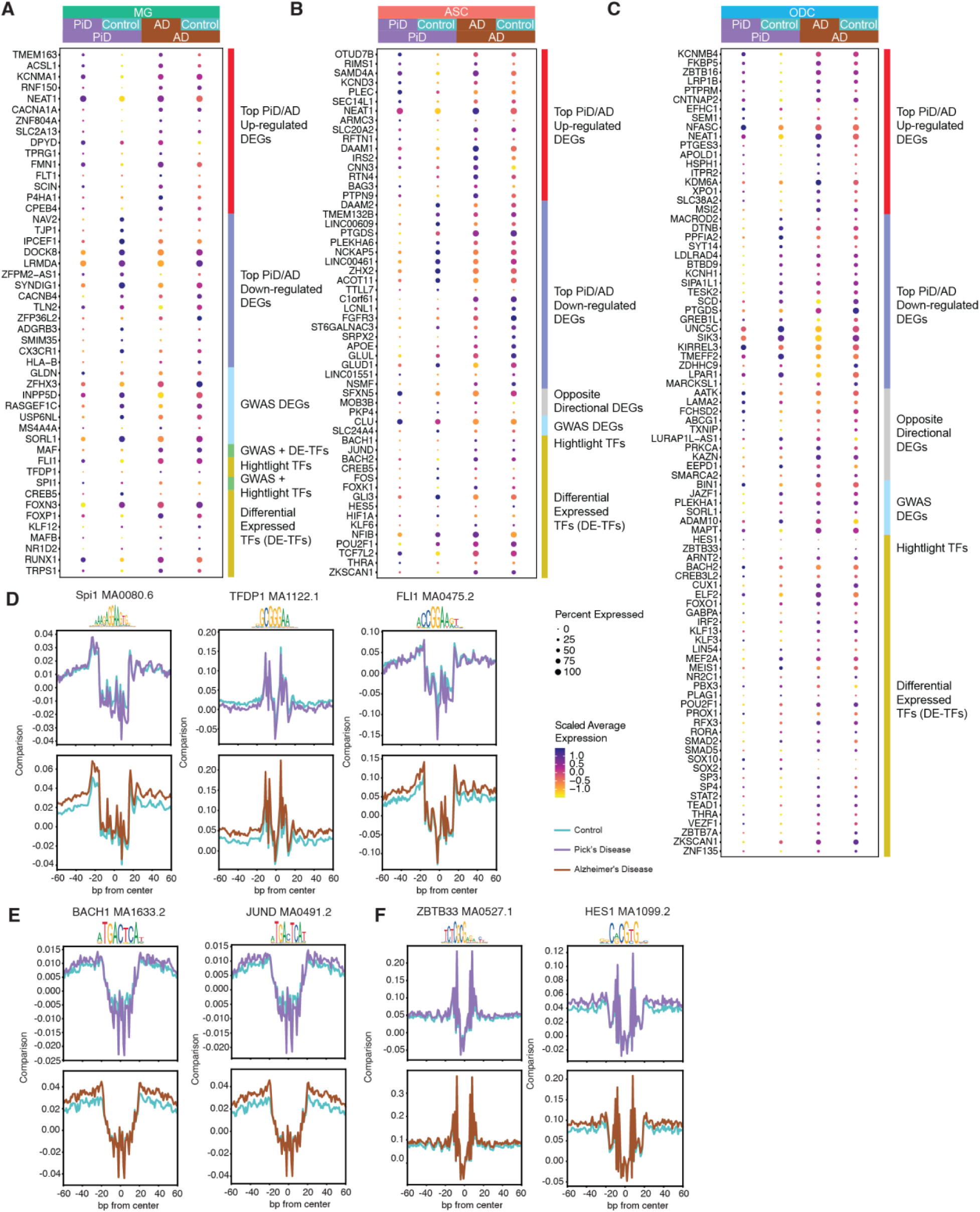
Aggregated footprints of TFs in MG, ASC, and ODC. (**A**, **B**, **C**) Dot-plot of the differentially expressed gene, differentially expressed GWAS risk genes, and TFs in PiD and AD versus their respective controls in MG (**A**), ASC (**B**), and ODC (**C**). (**D**) Aggregated TF footprints of *Spi1* (MA0080.6), *TFDP1* (MA1122.1) and *FLI1* (MA0475.2) in PiD and AD. (**E**) Aggregated TF footprints of *BACH1* (MA1633.2) and *JUND* (MA0491.2) in PiD and AD. (**F**) Aggregated TF footprints of *ZBTB33* (MA0527.1) and *HES1* (MA1099.2) in PiD and AD.

**Fig. S7.**
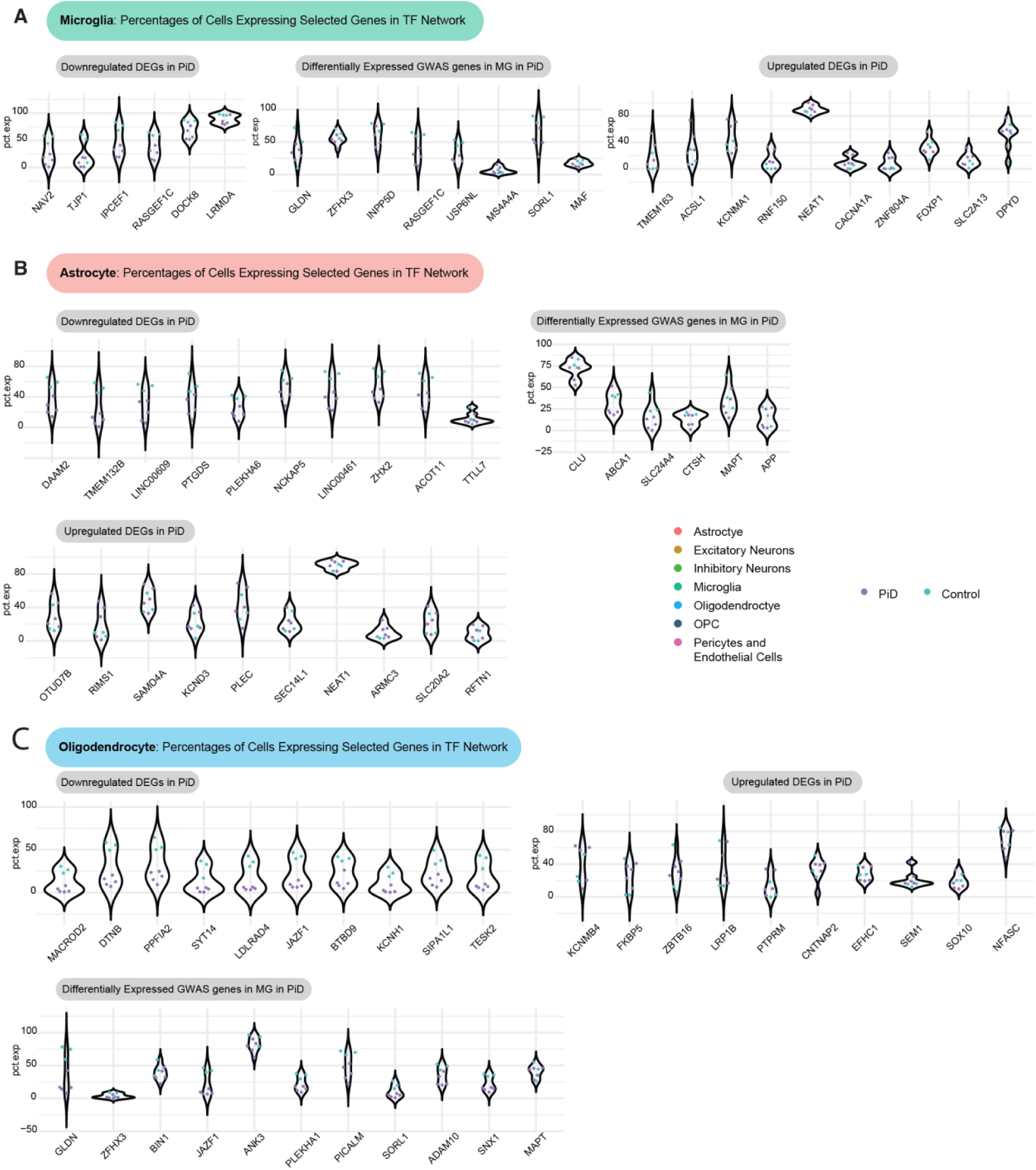
Percentages of cells expressing selected genes in TF. (**A**) Microglia: percentages of cells expressing selected genes in the TF network. (**B**) Astrocyte: percentages of cells expressing selected genes in the TF network. (**C**) Oligodendrocyte: percentages of cells expressing selected genes in the TF network.

**Fig. S8.**
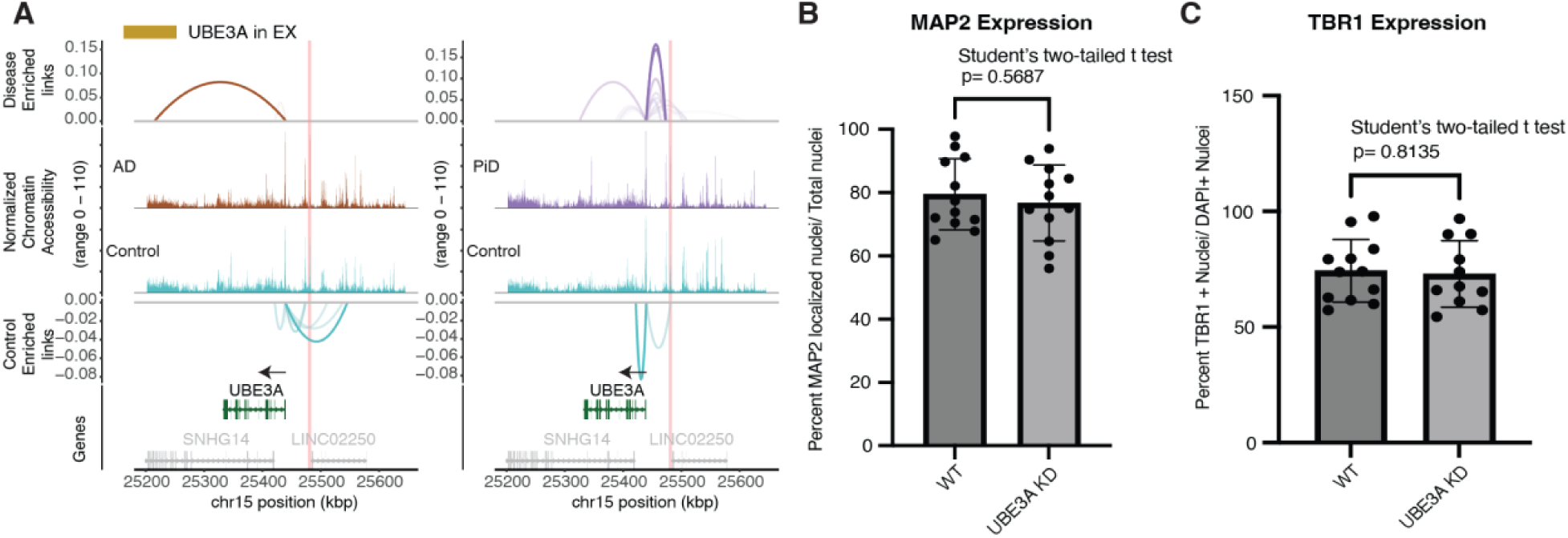
UBE3A enhancer accessibility and iPSC-derived neurons confirmation. (**A**) Delta co-accessibility of UBE3A and its open chromatin regions in EX for both AD and PiD with their corresponding controls. Highlighted regions in salmon represent CRISPR-edited enhancer regions to the UBE3A. (**B**) Quantification of DAPI nuclei colocalized with MAP2 expression, normalized to total nuclei. (**C**) Quantification of TBR1 positive nuclei normalized to total nuclei. Points represent individual images, n=4 per coverslip, n=1 coverslip per 3 differentiation replicates.

**Fig. S9.**
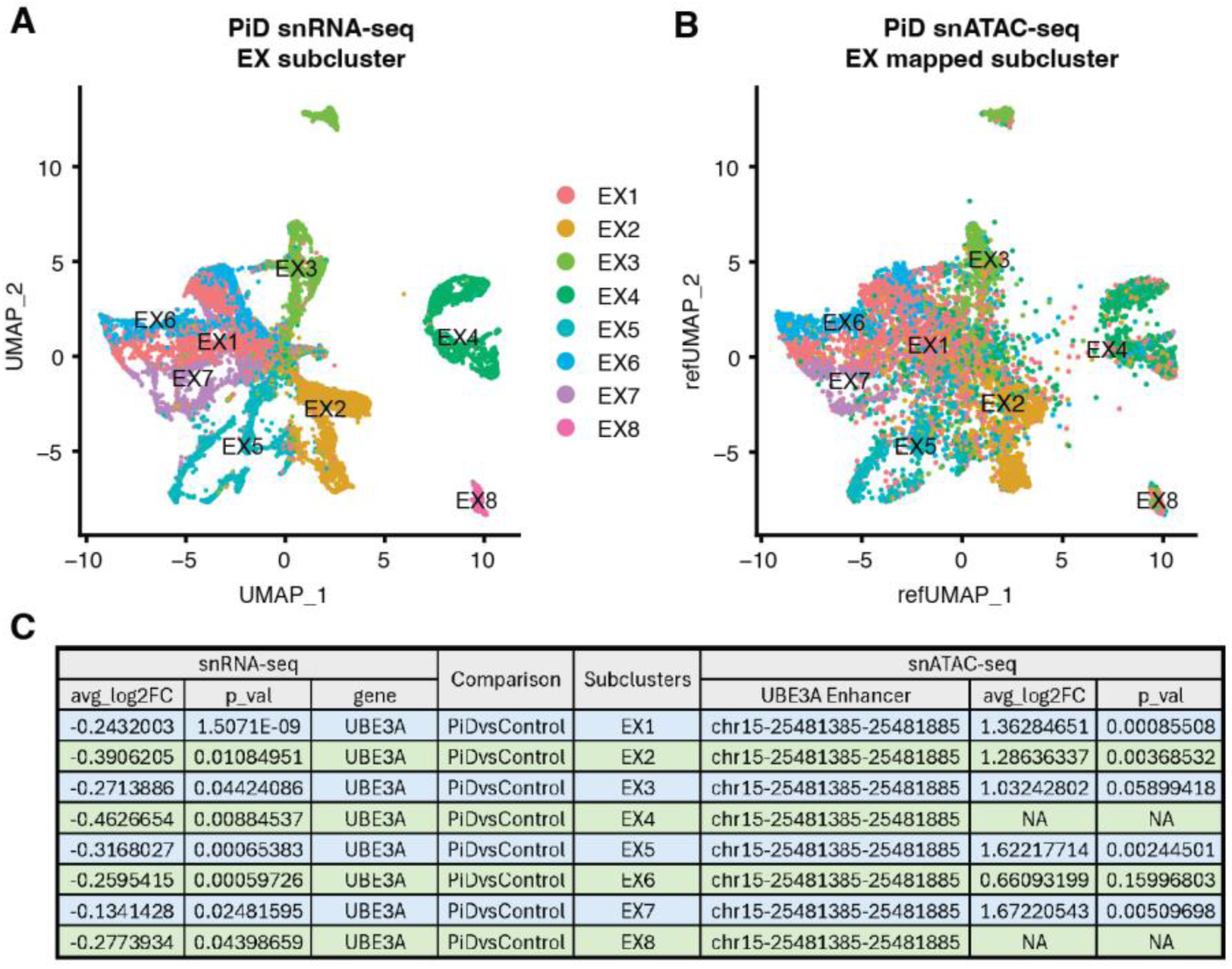
Subclustering analysis on EX. (**A**) UMAP of snRNA-seq EX subcluster. (**B**) UMAP of snATAC-seq EX subcluster. (**C**) Summary table of differential expression and accessibility analyses of EX subcluster.

**Fig. S10.**
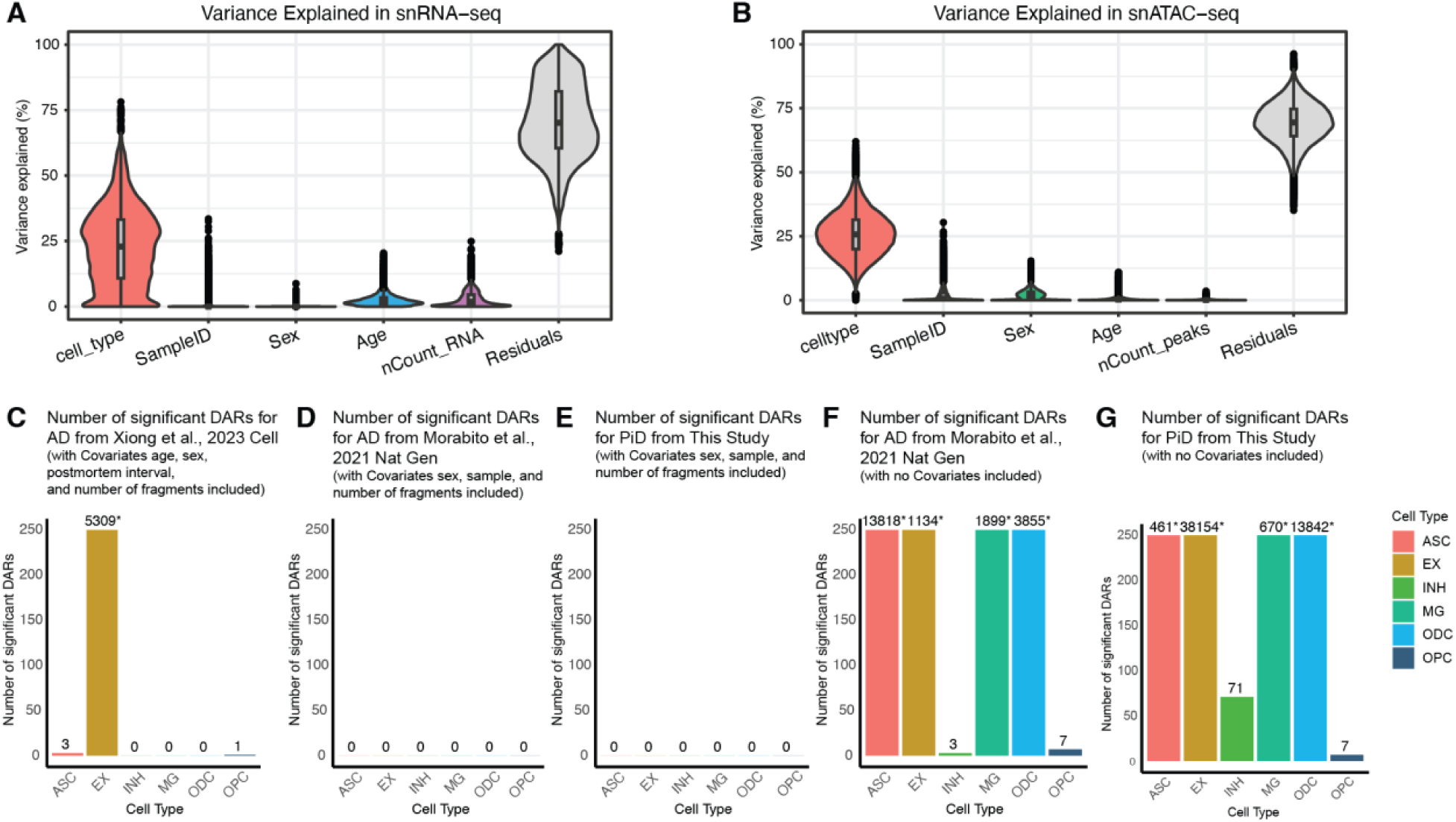
Covariate partitioning and impact of covariate selection on the number of statistically significant differentially accessible regions (DARs). (**A**) Source of variation analysis showing the proportion of gene expression variance explained by each covariate used in the differential expression model in snRNA-seq. (**B**) Source of variation analysis showing the proportion of chromatin accessibility variance explained by each covariate used in the differential expression model in snATAC-seq. (**C**) Number of statistically significant DARs per cell type with covariates (age, sex, postmortem interval, number of fragments) for AD data from Xiong et al., 2023, Cell (27). DARs were reprocessed using the same covariates as in snRNA-seq. (**D**) Number of statistically significant DARs per cell type with covariates (sex, sample, number of fragments) for AD data from Morabito et al., 2021, Nature Genetics (17). (**E**) Number of statistically significant DARs per cell type with covariates (sex, sample, number of fragments) for PiD data from this study. No covariates were used in the original analysis. (**F**) Number of statistically significant DARs per cell type without any covariates for AD data from Morabito et al., 2021, Nature Genetics (17). (**G**) Number of statistically significant DARs per cell type without any covariates for PiD data from this study.

**Table S1. Summary of supplementary tables providing metadata, marker analysis, and cell counts for PiD and AD datasets.**

(**Table S1A**) Metadata for PiD and AD. (**Table S1B**) Results from snATAC-seq FindAllMarkers analysis on gene activity. (**Table S1C**) Cell counts for each cell type across snATAC-seq and snRNA-seq datasets. (**Table S1D**) Results from snRNA-seq FindAllMarkers analysis on gene expression.

**Table S2. Summary of snATAC-seq peaks and differential accessibility regions (DARs) in PiD and AD datasets.**

(**Table S2A**) Complete peak set of snATAC-seq for both PiD and AD. (**Table S2B**) Summary counts of peak type and biotype in PiD and AD DARs (p < 0.05). (**Table S2C**) DAR analysis for PiD vs Control (pct > 0.05), including all statistics, not just significant ones.

**Table S3. Fine-mapping and GWAS enrichment analysis in PiD and AD datasets.**

(**Table S3A**) Overlap of snATAC-seq peaks with SuSiE fine-mapped credible sets (PIP > 0.95) and Xiong et al., 2023 (*27*). (**Table S3B**) Complete list of SuSiE fine-mapped credible sets (PIP > 0.95). (**Table S3C**) GWAS gene enrichment analysis in PiD DGE.

**Table S4. Differential gene expression (DGE) analysis in PiD and AD datasets.**

(**Table S4A**) DGE results using MAST glm for PiD vs Control. (**Table S4B**) DGE results using MAST glm for AD vs Control.

**Table S5. Human-gain enhancers (HGE) overlapped with snATAC-seq peaks in PiD and AD datasets.**

**Data S1. iPSC CRISPR KO Karyotype and Pluripotency Data Files.**

**(File 1)** Project_Report_UBE3A Swarup_6_30_22.pdf contains the design and results of the UBE3A CRISPR knockout experiment, including the knockout design strategy, experimental outcomes, quality control data for generated cell lines, and validation of iPSC characteristics. **(File 2)** Microarray REPORT CLG-46814_ADRC76.pdf is the microarray report for the parental iPSC line ADRC76, derived from fibroblasts and provided by the UCI Alzheimer’s Disease Research Center (ADRC) iPSC Core. **(File 3)** Microarray REPORT CLG-46815_C40.pdf corresponds to Clone 40 derived from the ADRC76 iPSC line. **(File 4)** Microarray REPORT CLG-46816_C14.pdf corresponds to Clone 14 derived from the ADRC76 iPSC line.

## Notes

https://datadryad.org/dataset/doi:10.5061/dryad.h9w0vt4t9

https://swaruplab.bio.uci.edu/scROAD/

https://www.ncbi.nlm.nih.gov/geo/query/acc.cgi?acc=GSE259298

### Competing Interest Statement

The authors have declared no competing interest.

### Summary of Updates

Several figures and the main text have been revised. Updates have been made to address the reviewer comments.

